# A phylogenomic perspective on interspecific competition

**DOI:** 10.1101/2023.05.11.540388

**Authors:** Nicolas Louw, Benjamin E. Wolfe, Lawrence H. Uricchio

## Abstract

Evolutionary processes may have substantial impacts on community assembly, but evidence for phylogenetic relatedness as a determinant of interspecific interaction strength remains mixed. In this perspective, we consider a possible role for discordance between gene trees and species trees in the interpretation of phylogenetic signal in studies of community ecology. Modern genomic data show that the evolutionary histories of many taxa are better described by a patchwork of histories that vary along the genome rather than a single species tree. If a subset of genomic loci harbor trait-related genetic variation, then the phylogeny at these loci may be more informative of interspecific trait differences than the genome background. We develop a simple method to detect loci harboring phylogenetic signal and demonstrate its application through a proof of principle analysis of *Penicillium* genomes and pairwise interaction strength. Our results show that phylogenetic signal that may be masked genome-wide could be detectable using phylogenomic techniques and may provide a window into the genetic basis for interspecific interactions.

**Data & code accessibility:** Data and code for this project are freely available in the repository linked below and will be permanently archived upon publication.

**Statement of authorship:** NL, BEW, and LHU designed the research; NL performed experiments; NL and LHU performed computational analyses; NL, BEW, and LHU wrote the manuscript.

**Code repository:** https://github.com/uricchio/ILSComp

## Introduction

Interactions between species are major determinants of the structure of ecological communities, and evolutionary processes are likely to shape the traits that determine interaction strength and outcomes. One widely-used empirical approach for characterizing the evolutionary basis of ecological interactions between closely related species compares phylogenetic distances to trait differences (Cavender-Bares et al., 2009). For each species pair in a community of interest, the evolutionary distance between the species can be regressed against some measure of the interaction strength between the species, such as a competition coefficient or relative growth rate. If competition plays a substantial role in limiting community membership, then species that co-occur naturally in biological communities might be less phylogenetically related than is expected under a null model (Webb et al., 2002). In contrast, if biotic interactions are weak determinants of community structure, environmental filtering could cause clustering of related species in space because only species with similar traits may be able to persist in a specific set of environmental conditions (Losos, 1996). In either case, evolutionary history should be informative of community composition and/or the strength of interspecific interactions, a pattern sometimes called “phylogenetic signal”.

Many empirical studies have now performed such analyses, sometimes finding that relatedness is a strong predictor of interspecific competitive differences (*e*.*g*., Jiang et al. 2010; Peay et al. 2012; Godoy et al. 2014) and sometimes not (*e*.*g*., Cahill Jr et al. 2008; Narwani et al. 2013; Naughton et al. 2015). The inconsistency of phylogenetic signal for competitive interaction strength across studies has been discussed and debated extensively, with potential explanations including contrasting effects of relatedness on niche and fitness differences (Mayfield and Levine, 2010) and differences in the phylogenetic or spatial scales considered (Parmentier et al., 2014; Graham et al., 2018; Jarzyna et al., 2021). Though this debate continues, most ecologists seem to agree that phylogenetic structure would be more likely to emerge for traits that determine species interactions than for traits that are unrelated to community assembly processes or outcomes (Mayfield and Levine, 2010).

The inference of phylogenetic trees, and more broadly the evolutionary processes that determine phylogenetic structure, could also play an important role in the detection and origination of phylogenetic signal. It is now well-appreciated amongst evolutionary geneticists that the genomes of many species are not well-represented by a single phylogeny – different portions of the genome can have different evolutionary histories (*e*.*g*., Hime et al. 2021; Meleshko et al. 2021). This raises the possibility that while the species level phylogeny may not always be suggestive of phylogenetic signal, subsets of the genome could have phylogenies that are strongly related to trait differences and/or processes determining community assembly. When a single phylogeny is inferred based on concatenated genetic or genomic data for a particular set of focal species, regressing trait differences on phylogenetic differences could provide evidence for or against phylogenetic signal, but provides no additional insight into the specific genomic loci that determine trait differences because the inferred evolutionary history is shared across all loci. Taking a phylogenomic approach, in which the species tree is replaced with the phylogeny at each genomic locus, might provide new insights into the evolutionary processes and genomic variation involved in community assembly processes.

When a portion of the genome has an evolutionary history that differs topologically from the genome background, this phenomenon is known as “discordance” between the local gene tree and the species tree (Degnan and Rosenberg 2009; see Box 1). The gene tree represents the phylogeny as inferred only from the genetic variants falling within a single locus in the genome (often the sequence of a gene, but possibly a contiguous segment of DNA that does not code for a gene). Discordance can arise through various evolutionary processes, such as introgression, gene duplication, and incomplete lineage sorting. For example, if hybridization occurs between two distinct species then the genomes of the hybrids become a patchwork of evolutionary histories characterized by the distinct histories of the parent species. Gene flow that results in the retention of genetic material from another species is known as “introgression”, and if the new genetic material confers a fitness advantage then the process is called “adaptive introgression” (Edelman and Mallet, 2021). Discordance can also arise through neutral processes and without any gene flow between species. For example, genome-wide genetic divergence suggests that humans are most closely related to chimpanzees and bonobos, but a substantial portion of our genomes are more closely related to gorillas than chimpanzees (Ebersberger et al., 2007). This pattern is expected when the internal branches of the species tree are short, and arises due to a process known as “incomplete lineage sorting” (Degnan and Rosenberg 2009; see Box 1).

Supposing that only a small subset of genes in the genome may be relevant to determining traits that affect co-occurrence patterns, phylogenetic signal may be more likely to arise at loci that are causal for trait differences than at the level of the genome background, especially when a gene harboring variation that is causal for the trait of interest has an evolutionary history that is discordant with the species tree. One recently proposed method known as `PhyloGWAS` leverages patterns of discordance to detect associations between traits and genomic loci (Pease et al., 2016; Wu et al., 2018), providing a window into genes that may be involved in causal differences in traits across species. However, `PhyloGWAS` has not (to our knowledge) been applied to interspecific interactions, perhaps in part because it was designed primarily for categorical traits.

Here, we consider the potential for phylogenomic approaches to “rescue” phylogenetic signal that is obscured at the level of the species tree and reveal genomic loci that may underly traits involved in community assembly. We develop a simple phylogenomic approach for detecting associations between gene trees and quantitative traits and apply it to experimental data from eight co-occurring fungal species from the *Penicillium* genus as a proof of principle. We show that our approach identifies genomic loci that are associated with competition strength in *Penicillium*, restoring phylogenetic signal that would not be detected with the species tree alone. Using gene annotation data, we show how such analyses can be used to generate new hypotheses about the ecological drivers of trait differentiation when such annotations are available. We discuss the interpretation of phylogenetic signal in studies of community structure in light of discordance patterns. Lastly, we propose ways to extend and adapt our approach to phylogenomic analysis of quantitative traits.

### Conceptual overview

The genomes of many species are composed of a patchwork of evolutionary histories. A pair of trees describing the evolutionary histories of different genomic loci are described to as “discordant” when the topology of these trees differ (Box 1). Since gene trees can be discordant, different regions of the genome may have different levels of evidence for association with quantitative traits as determined by the correlation between evolutionary distances and trait differences. A phylogenomic approach can leverage discordance to identify regions of the genome with particularly strong associations with traits of interest.

In its most basic form, our approach can be thought of as an outlier scan for genomic loci that are strongly correlated with trait differences between species. As with all outlier scans, it is critical to differentiate between expected variation under a reasonable null hypothesis and variation that is in excess of the correlations expected under the null. We discuss the role of various evolutionary processes in generating signals of association between gene trees and traits, describe the basic mechanics of an example analysis in the following section, and then turn to the generation of a null distribution that allows us to determine statistical significance.

### Evolutionary processes and the detection of phylogenomic associations

Several distinct evolutionary processes can lead to discordance between gene trees and species trees (Box 1), and hence these processes determine the statistical performance of phylogenomic association approaches (Box 2). As an example, we visualize discordance for a sample dataset of eight *Penicillium* species by plotting a sample of gene trees (blue) along with the consensus phylogeny (dark gray) in Fig. 1. When discordance is low, most of the blue trees will have identical topologies to the species tree. When discordance is very high, most gene trees will have different topologies than the species tree. Our example displays high discordance amongst the closely related clade (inluding *P. mb, P. polonicum, P. verrucosum, P. solitum*, and *P. bioforme*) and more moderate discordance amongst the three more diverged species. Note that discordance can arise due to tree inference errors or ortholog misidentification, but is also expected due to evolutionary processes such as incomplete lineage sorting and introgression (Box 1).

**Figure 1:**
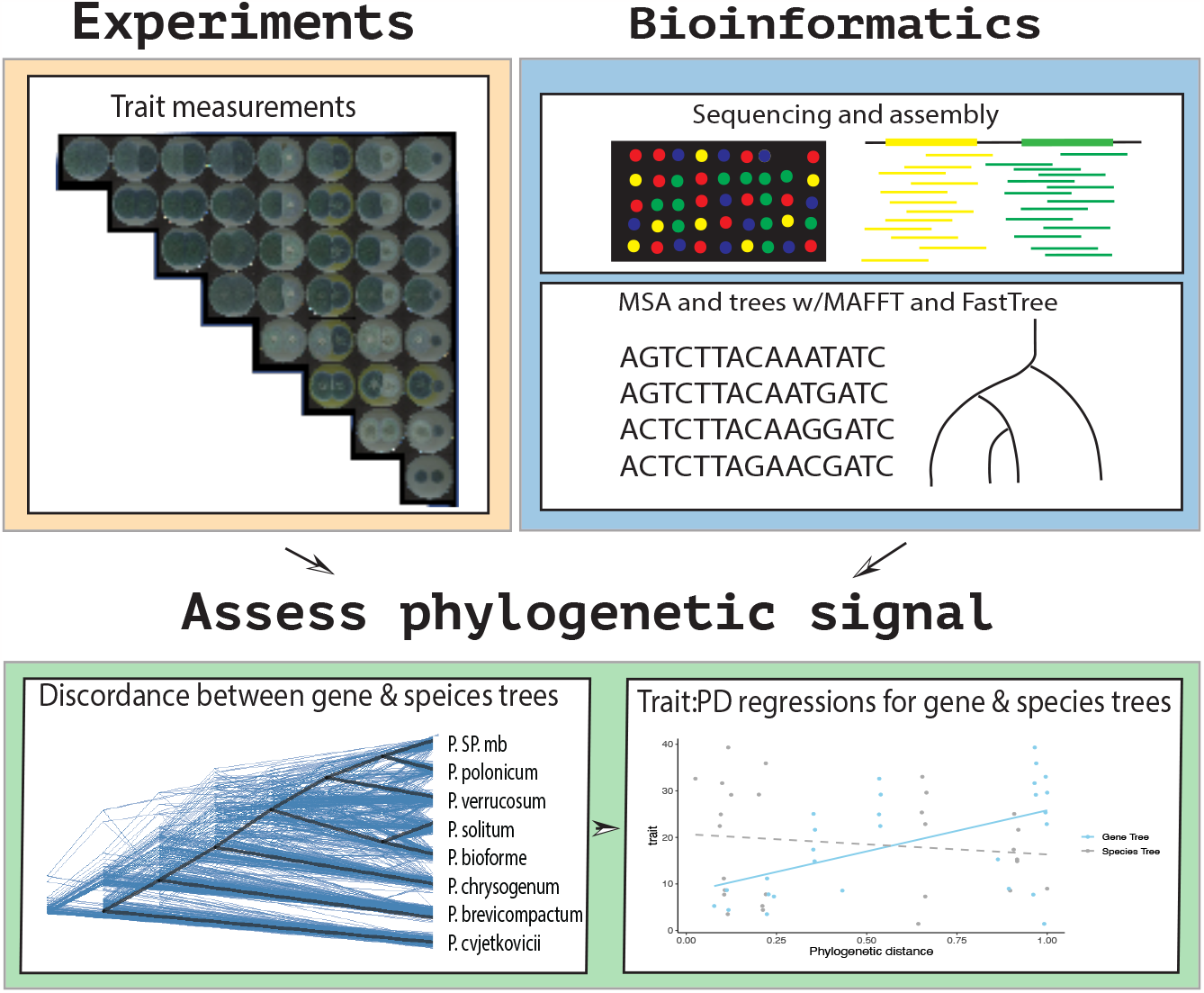
Workflow for our proof of principle analysis. We measured competition strength using pairwise growth experiments (upper left), but any quantitative trait can be analyzed within our framework. We used whole genome sequences of the eight species in our analysis to infer orthologs, make alignments and infer gene trees (upper right). We assessed phylogenetic signal by comparing evolutionary differences using gene trees with trait differences (lower panel). The tree on the bottom left illustrates discordance between our species tree and subset of the gene trees. The tree figure was made with `ggtree` (Xu et al., 2022). Note that the regressions shown on the lower right hand side are hypothetical data illustrating a positive result for a particular gene tree (blue) and no correlation for the species tree (gray).

When discordance is very low (as is expected when the internal branches of the species tree are very long and there is relatively little evidence for hybridization), most gene trees will have branch lengths and topologies that are very similar to the species tree. By definition, this will make the identification of associated genes quite difficult using our approach (*i*.*e*., gene trees at both causal genes and non-causal genes will have very similar correlation coefficients with trait values). However, this also means that relatively small distortions in topology of the gene tree (relative to the species tree) can be detected in this limit, because such topologies may be very unlikely under a reasonable null model. Moreover, gene trees that have have branches that have accumulated very large numbers of substitutions due to selection for trait differentiation may also have stronger correlations with trait values than the genome background, even if the topologies are identical.

In contrast, taxa with a large amount of discordance (as is expected when the internal branches of the tree are very short) by definition have many genes that are discordant from the species tree. If most genes have topologies that are not identical to the species tree and a small portion of genes explain most of the variance in interspecific trait values then these individual genes are more likely to display phylogenetic signal than the species tree. If instead there are many causal genes and each explains only a small portion of the variance in interspecific trait differences, then each individual gene may have very weak associations but the species tree – which represents a consensus over gene trees – may be more likely to be correlated with the trait. In sum, both high discordance and low discordance taxa could be analyzed in our framework, but the evolutionary processes underlying potentially significant loci differ between these regimes and are likely to have substantial impacts on the statistical performance of the approach (Box 2).

### Example workflow

Our approach scans the genome for genes with gene tree topologies and branch lengths that are more strongly correlated with trait differences than the genome background (Box 1; Fig. 1). The analysis is composed of several steps. First, a set of species of interest is identified and trait data are collected. For the proof-of-principle analysis in this Perspective, we performed pairwise competition assays for eight *Penicillium* species (Fig. 1, upper left), but any quantitative trait related to interspecific competition (or any other ecological process) could be analyzed in a phylogenomic framework such as the one proposed here. For example, commonly collected datasets including pairwise competition estimates between plant species (*e*.*g*., Cahill Jr et al. 2008) or body sizes of mammals (*e*.*g*., Davies et al. 2012), both of which have been hypothesized to be related to co-occurrence patterns, could easily be studied in our framework. When trait data are naturally pairwise and directional (*e*.*g*., competition coefficients under a Beverton-Holt model; Brännström and Sumpter 2005) the pairwise estimates can be used directly, while if the trait data naturally correspond to individual species (*e*.*g*. body sizes) then some measure of the difference in values between species can be used.

We then perform bioinformatic steps that are standard to many genomic research workflows – we assemble genomes, identify orthologous genes, build alignments of the orthologs, and infer a phylogeny for each set of orhtologs (Fig. 1, upper right). In our example analysis, we used `OrthoFinder` to identify orthologs (Emms and Kelly, 2019) and `MAFFT` (Katoh and Standley, 2013) to build alignments, while gene trees were inferred with `FastTree` (Price et al., 2010) and the species tree was inferred with `RAxML-NG` (Kozlov et al., 2019), but numerous other tools could be used for each step and might have advantages that are specific to particular use cases (*e*.*g*., Maher and Hernandez 2015). We include only single-copy orthologs in our analysis (*i*.*e*., genes for which each species has a single copy of the gene in its genome) since it is much more straightforward to infer and interpret phylogenies for this set of genes, but we consider extensions that relax this constraint in the Discussion section.

The final step is to assess phylogenetic signal by comparing phylogenetic distances (PD) to trait values (Fig. 1, bottom right). For each gene and each focal species, we compute the naive correlation coefficient between the pairwise competition coefficients and phylogenetic distance to each of its competitors. This provides a distribution over correlation coefficients across the genome for each species, and our goal is to identify genes with especially strong correlations. If we were interested only in obtaining a list of genes with the strongest correlations, we could stop at this step and simply list the genes with the strongest correlations. However, all distributions have tails, and an essential part of our approach is to determine a reasonable null distribution against which we can assess the significance of single gene correlations with trait values. We turn to this problem in the next section.

### Null model

To assess statistical significance, our method relies on the generation of a null distribution of correlation coefficients between phylogenetic distances and pairwise interspecific trait differences under a Multi-Species Coalescent model (Edwards et al., 2016). The key idea behind this approach is that gene trees are contingent on a species tree – they cannot evolve in a way that is completely unconstrained, but rather according to an evolutionary model that describes the distribution of gene trees given the species tree (Box 1). Whereas many prior approaches designed to assess the evidence for phylogenetic signal have sought to use the covariance structure imposed by a species tree to effectively remove co-dependence between observed trait values (Felsenstein, 1985; Münkemüller et al., 2012), our phylogenomic approach seeks to directly account for this co-dependence in the null model.

Since the species tree constrains the evolution of the gene trees, our approach requires a high quality consensus species tree inferred from a set of genomic loci. Given this species tree, we can use simulations under the multi-species coalescent model to approximate the expected distribution of correlation coefficients under the null. Genomic loci that have associations in excess of a threshold determined by a multiple testing correction can be considered significantly associated. Even in the absence of statistical significance at specific individual loci, the set of genome-wide gene trees could reveal phylogenetic signal that was not detectable at the whole genome/species tree level. Methodological details of each of these steps are provided in the Supporting Information, but we stress that we have opted for simplicity in this Perspective as a proof of principle. Many of the choices we made in this study could be optimized to improve power or decrease false positive rates and detect specific evolutionary patterns we have not considered, as elaborated upon in the Discussion.

#### Box 1

**Evolutionary processes and discordance**

Discordance describes the phenomenon in which different portions of the genome may be described by different evolutionary histories. A gene tree (typically inferred using variation at a single locus in the genome) is discordant with the species tree (typically inferred using whole-genome data or a susbstantial portion of the genome) if the two trees have different topologies. Discordance can arise through a number of distinct evolutionary processes, including introgression, incomplete lineage sorting, and gene duplication.

##### Introgression

Introgression occurs when two species interbreed, resulting in gene flow into one (or both) of the parent species. Introgressed genetic elements can result in the acquisition of novel traits, or can provide a fast route to local adaptation (Edelman and Mallet, 2021), though introgression does not have to be adaptive and may even be costly in some cases (Harris and Nielsen, 2016). Introgression also causes the genome to become a mosaic of evolutionary histories, represented by the distinct histories of the parent species, and such introgression events are detectable through a number of computational methods (Hibbins and Hahn, 2022). Genes introduced through introgression could therefore be both important contributors to functionally important trait differences across species, and could result in patterns of discordance between the introduced locus and the species phylogeny.

##### Incomplete lineage sorting

Discordance can arise even when there is no exchange of genetic material between species. For example, consider the evolutionary history of a gene tree (colored lines), which is constrained by the species tree (outer black lines). The common ancestor to species *B* and *C* at this locus must have lived at some time prior to the split between these species in the species tree. Common ancestry need not be shared immediately upon the speciation event for any given locus – the common ancestor could be an individual who lived at any time in the past preceding this speciation event. If the common ancestor to *B* and *C* is an individual who lived prior to the speciation event between *A* and (*B, C*), then there is a chance that the topology of the gene tree will differ from the species tree. This phenomenon is known as “incomplete lineage sorting”, because the lineages do not “sort” on the gene tree in the same order as the species tree. A mathematical framework that describes the likelihood of ILS is known as the Multi-Species Coalescent (MSC) model (Degnan and Rosenberg, 2009; Edwards et al., 2016). Under the model, the species tree is a container that constrains the distribution of topologies of gene trees. ILS is more likely when the internal branches of the species tree are short relative to population size. If the internal branches of the species tree are long, then most gene trees are expected to have the same topology as the species tree. Note that the classic version of the MSC does not allow for introgression/hybridization events, but they can be incorporated through simulation.

##### Gene duplication and ortholog detection

Gene duplication events can also lead to apparent discordant histories (Galtier and Daubin, 2008). In a gene duplication, a DNA sequence is duplicated during replication, resulting in multiple copies of a gene. Such duplication events can make it more difficult to properly infer orthologs (*i*.*e*., sequences arising from a common ancestor). For example, if a set of putative orthologs actually contains a mixture of genes arising from the two duplicated gene copies, then the underlying topology may differ from the species tree. For this reason, empirical studies of discordance typically rely on a set of high confidence single-copy orthologs and/or take great care in mapping putative duplication events.

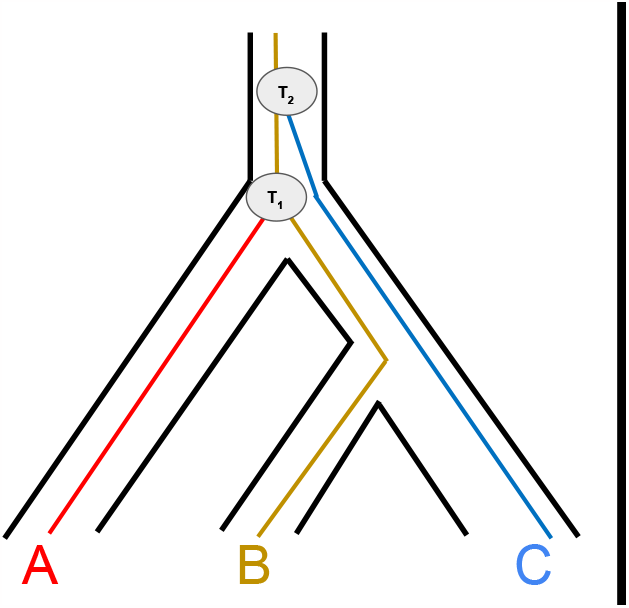

The dark black lines represent the species tree, while the colored lines represent a gene tree. The species tree and gene tree are discordant because B and C are the most closely related in the species tree but not the gene tree. *T*_1_ indicates the time when the lineage from species *A* and *B* coalesce, whereas *T*_2_ repersents coalescence of (*A,B*) with *C*.

Our approach is conceptually related to `PhyloGWAS` (Pease et al., 2016), which scans the genome for genes that have larger than expected numbers of nonsynonymous substitutions that perfectly correlate with a categorical trait of interest. If the trait of interest is dispersed across the species tree (rather than being concentrated into a single monophyletic clade) then correlations between substitutions and traits could be due to 1) incomplete lineage sorting, in which a the species that share the trait are monophyletic in the gene tree but not the species tree, or 2) recurrent substitutions on distinct parts of the tree, or 3) introgression. `PhyloGWAS` assesses the likelihood that the substitutions arose due to incomplete lineage sorting (which is taken as a neutral null model) by simulating gene trees under the multi-species coalescent model. If a gene has a larger than expected number of correlated nonsynonymous substitutions such that the null hypothesis of incomplete lineage sorting can be rejected based on the simulated distribution of substitutions, then the gene is considered to be a candidate locus for the determination of the trait.

#### Box 2

**When is discordance informative for studying trait differences?**

##### Discordance can be biologically informative

Under a neutral coalescent model, genes with discordant histories are no more likely to have functional relevance to community assembly or other ecological processes than genes that have similar topologies to the species tree. However, if traits evolve neutrally and are determined by genetic variants within a subset of genes in the genome, then under most evolutionary models the number of genetic differences in this subset of causal genes should be predictive of trait differences between species. If the causal genes have topologies that are distinct from the genome background, then they may also be more correlated with trait differences than average genes in the genome. Hence, in the high discordance limit, genes that are correlated with trait differences in excess of the amount that is expected due to the phylogenetic structure of the species tree could be determinants of trait differences. In contrast, if causal genes have the same topology as the species tree – as would be expected in the low discordance limit – then phylogenetic signal should be detectable at the species tree level when it exists, but it may be impossible to detect correlations with the causal genes using discordance signals.

##### The role of non-neutral evolutionary processes

Natural selection could also play a role in the determination of trait differences, and indeed a role for selection may be likely for traits underlying community or population outcomes. If selection determines trait variation, then genes that cause trait differences may have excessive levels of differentiation along selected branches. In the context of community assembly, organisms that evolved to occupy similar niches might have more shared variation in ecologically relevant genes than is expected due to their overall relatedness. Selection could therefore affect the inference of evolutionary distances within causal genes, potentially making them easier to detect relative to the genome background.

##### Empirical distribution of discordance

The amount of discordance between gene trees and species trees can be summarized using a number of different statistics, such as the Robinson-Foulds (RF) metric (Robinson and Foulds, 1981). When discordance is low, many gene trees will have the same topology as the species tree and hence an RF value of 0 relative to the species tree, while increasing discordance results in a broader distribution of RF values. Phylogenomic association methods may be most applicable in the high discordance limit – by calculating RF distributions across sets of gene trees (or other summaries of discordance) researchers can therefore determine the suitability of such approaches for their datasets. However, this does not preclude the possibility that a small subset of genes may have evolutionary histories that are highly unlikely given the phylogeny in the low discordance limit. For example, genes with a causal role in trait differentiation that are introduced via introgression should be easier to detect in the low discordance limit, because their topologies and correlations with trait values may be very unlikely under a neutral coalescent model.

A substitution that is shared only by the subset of species that have a particular trait value induces a split (or bipartition) among the species, *i*.*e*. it separates all of the species in the phylogeny into two groups. Since `PhyloGWAS` scans the genomes for genes that have a larger-than-expected number of such substitutions, it is analogous to searching for gene trees that contain a longer-than-expected branch corresponding to this bipartition. By regressing the gene tree branch lengths on the traits (rather than counting substitutions along the branch of interest) we can naturally extend this idea to quantitative traits without determining an *a priori* split between the species. Effectively, we are searching for gene trees where the number of substitutions along the branch is more strongly predictive of differences in trait values than we would expect given the species-level phylogeny. This is a desirable property for quantitative traits because there is no natural grouping of the species, as there is with a categorical trait.

## Results

### Assessing phylogenetic signal under a multi-species coalescent model

We first considered the possibility of assessing the significance of correlations between gene trees and traits by performing naive regressions, without any correction for the species tree. We hypothesized that this approach would work poorly in both the low and high discordance limit. To test this, we simulated a neutrally evolving trait, the value of which on each of the leaf nodes of the tree is determined by substitutions that occur along each branch of a single causal gene tree. Then we regress trait differences against phylogenetic distances for each causal gene tree without any correction for the species tree. The gene trees were simulated according to the multi-species coalescent model, given a species tree for the 8 *Penicillium* species in our study (see Supporting Information). For each causal gene and trait distribution, we also simulated 1,000 additional non-causal genes sampled on the same species tree (Fig. 1A). We repeated this procedure 1,000 times, regressing the phylogenetic distances determined by gene tree branch lengths on squared differences in trait values for the species at the leaf nodes for both the true causal gene and the 1,000 non-causal genes. If this method were effective, we would hope that causal-genes tend to have lower p-values than non-causal genes, and that the distribution of p-values under the null would be approximately uniform.

Results of these regressions (summarized by *p*-values) are reported in Fig. 2. When discordance between gene trees and species trees is very high, the true causal gene is often more correlated with trait values than a random gene sampled from the null distribution of gene trees (Fig. 2A), suggesting that under high discordance a naive calculation of correlation coefficients would have some power to identify causal loci under this simple model. However, as the amount of discordance decreases (*i*.*e*., as population sizes get smaller relative to branch lengths of the species tree), the distribution of *p*-values under the null looks increasingly similar to the distribution of *p*-values for causal gene (Fig. 2B-C). This is expected, because branch lengths and topologies are increasingly similar between gene trees and species trees when discordance is low. Low discordance results in severely inflated *p*-values for correlations between gene tree distances and trait differences (Fig. 2D) and a ROC curve suggests no ability to distinguish causal loci from the background (Fig. 2E). In contrast, high discordance results in inflation of *p*-values only under about *p* = 1*e −* 2 and provides some power to distinguish causal and non-causal genes (Fig. 2D-E). Note that while this qualitative pattern is expected with any species tree, the distribution of *p*-values for both null and true positive genes depends on the topology of the tree and details of the evolutionary model that generates trait differences across species.

**Figure 2:**
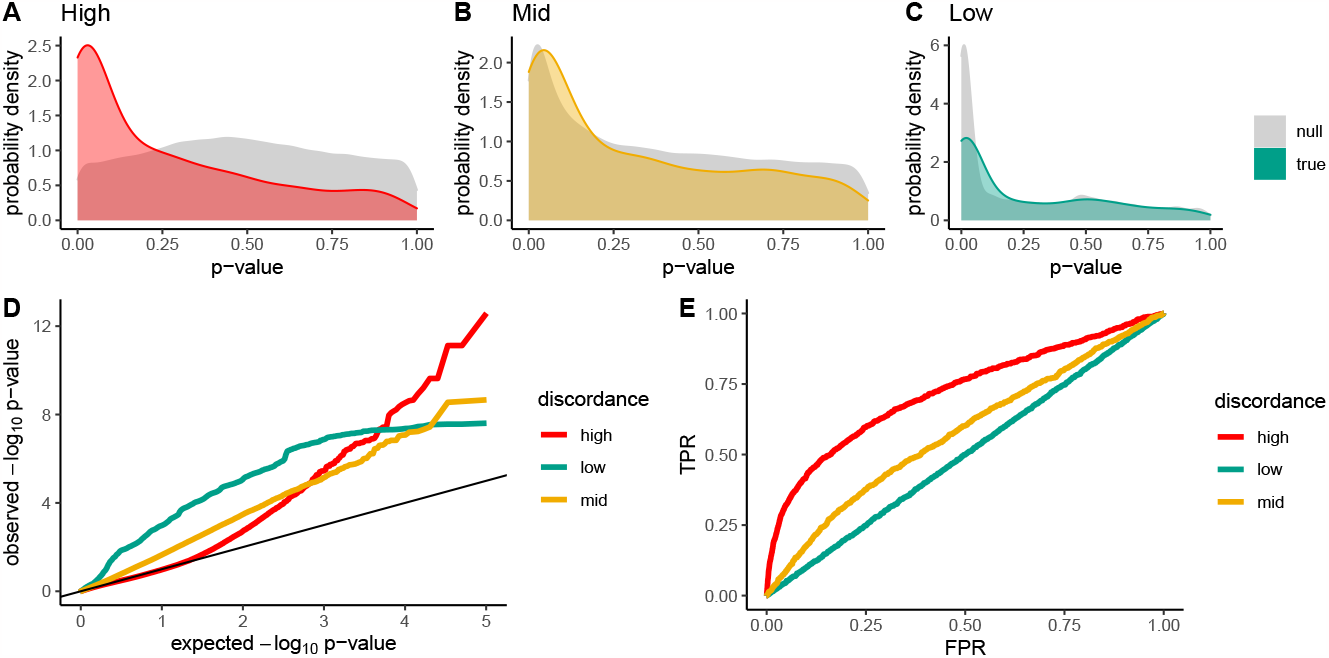
Statistical performance of naive correlations between gene trees and trait values simulated under the MSC model. A-C: Probability density of *p*-values obtained for the correlation between phylogenetic distances and simulated trait differences in trait values under the “null” (*i*.*e*., random gene trees) and for the “true” causal gene tree for each of three levels of discordance. Each panel in this figure takes *P. bioforme* as the focal species and the plotted distributions correspond to the marginal distribution of *p*-values across all seven competitor species – each focal species has it’s own such null distribution, and we choose *P. bioforme* as a representative example. D: QQ-plot for *p*-values obtained under the null for high, low, and moderate gene/species discordance. E: ROC curve for identification of causal genes based on *p*-values of the correlation between gene tree branch lengths and pairwise trait differences.

While each of these computational experiments resulted in some degree of inflation of *p*-values under the null, it is clear that lower discordance results in more inflation. We therefore repeated this computational experiment, but using phylogenetic least squares (PGLS) as implemented in `ape` and `geiger` (Paradis and Schliep, 2019; Pennell et al., 2014) as an attempt to control for phylogenetic structure. PGLS is similar to the classic method of phylogenetically independent contrasts in that it allows us to account for the expected trait covariance structure induced by the species tree (PICs; Felsenstein 1985), but is more flexible and allows for the estimation of additional parameters (Rohlf, 2001).

As expected, this had little effect on the analysis when discordance is high – under high discordance, the topology of gene trees is only very weakly affected by the structure of the species tree, so including the species tree does not correctly account for the co-variance structure in the data (Fig. S1). PGLS did moderately decrease inflation of *p*-values under *p* = 1*e −* 1 for low and moderate discordance (Fig. S1D), but did not fully correct the *p*-value distribution under the null. ROC curves were similar between PGLS (Fig. S1E) and naive correlation analyses (Fig. 2E).

### A model-based correction accounting for phylogenetic relatedness

We next used the Multi-Species Coalescent (MSC) model to determine a null distribution over correlation coefficients between trait values and gene tree branch lengths, given a species tree and a trait distribution at the tips of the tree. As in `PhyloGWAS` (Pease et al., 2016; Wu et al., 2018), we use a simulation-based approach because calculations under the MSC are not straightforward and depend on both tree topologies and trait distributions that are not known *a priori*. We simulate gene trees and calculate the null distribution over correlation coefficients, which we can then use to identify loci with unexpectedly large correlations. We set the population size using an empirically driven approach, which minimizes the difference between the observed and simulated distributions of correlation coefficients to infer an appropriate population size (see Fig. S3).

In Fig. 3, we plot both the observed distribution of correlation coefficients and the distribution determined under the MSC model for all 8 species of *Penicillium* in our experiments, where the trait values correspond to RCC. In general these simulation-based null distributions capture the overall shape and location of the observed distributions. For two species, (*P. brevicompactum* and *P. cvjetkovicii*, Fig. 3A and Fig. 3D, respectively) the observed distributions were peaked in similar locations as the MSC-based distributions, but were considerably wider. One peak expected under the MSC (Fig. 3D) is visibly absent in the observed data. Nonetheless, the breadth of the simulated null distributions is generally similar to the observed distributions, suggesting that the null distributions are useful for the calculation of simulation-based *p*-values.

**Figure 3:**
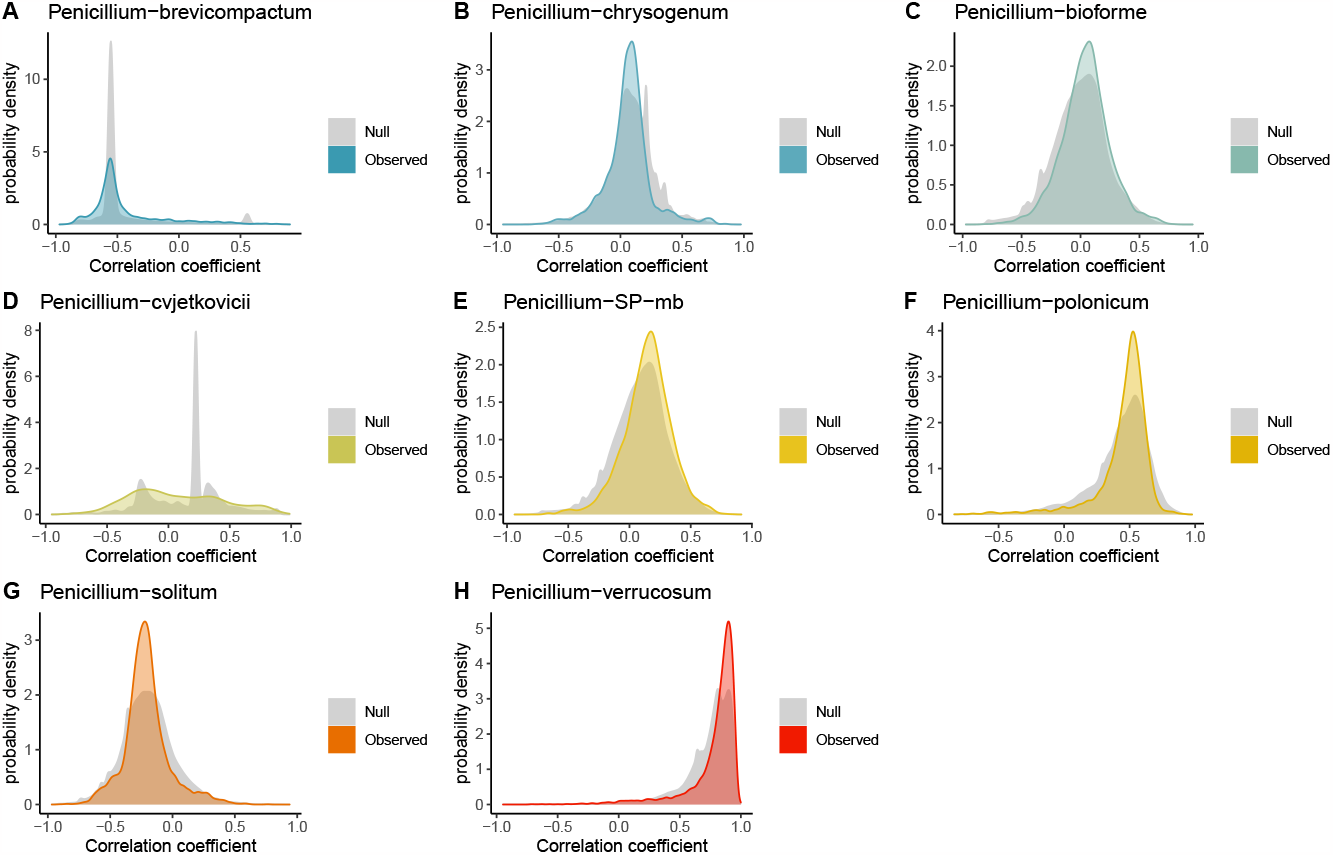
Comparison between simulated null and observed distributions of correlation coefficients across gene trees for each focal species. A-H: MSC-based null distribution of correlation coefficients between RCC and phylogenetic distances (gray) as compared to the observed distribution obtained from the collection of inferred gene trees (colors), for each focal species of *Penicillium* in our study.

We then compared the distribution of *p*-values obtained under our simulation-based approach to *p*-values obtained through naive correlations between gene tree distances and trait values (Fig. 4). We corrected *p*-values by using the distribution of correlation coefficients obtained under the MSC model as a null – the corrected *p*-value is the proportion of simulations in which the correlation coefficient exceeds the observed correlation coefficient. Naive calculations of correlation coefficients result in distributions of *p*-values that can be both strongly deflated and inflated, depending on how the species tree constrains gene trees for each individual species (Fig. 4A). Corrected *p*-values were approximately uniformly distributed, but may slightly over-correct for some species (*e*.*g*., *P. verrucosum*) or under-correct for others (*e*.*g*., *P. brevicompactum*). Note that in general, statistical performance of our approach will depend on aspects of the evolutionary model that are not known *a priori*. In Fig. S2 we display results of simulated traits with polygenic architecture (multiple causal genes) in the high discordance limit. As the number of causal loci increases, there is a sharp penalty to statistical power. However, sampling of more species increases power when discordance is high.

**Figure 4:**
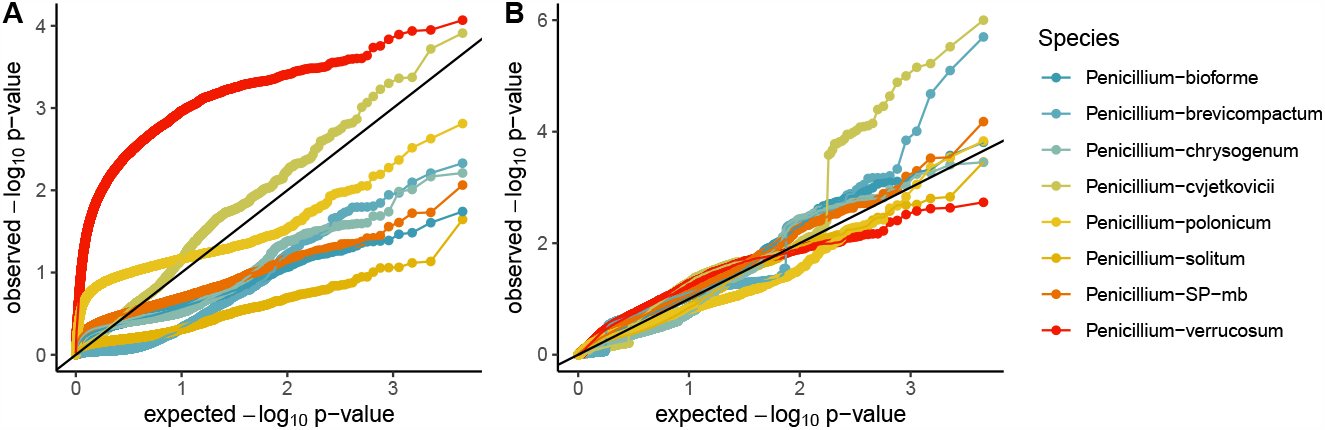
Comparison of p-value distributions both with and without a phylogenetic correction. A: QQ-plot for *p*-values obtained by naive correlation analysis between RCC and phylogenetic distances, for each focal species of *Penicillium*. B: QQ-plot for *p*-values obtained under our simulation-based null model between RCC and phylogenetic distances, for each focal species of *Penicillium*.

### Genomic loci associated with the strength of competition in *Penicillium*

Given our model-based null distributions of correlations between phylogenetic distances and trait differences, we scanned for outliers with correlations that were higher than expected under the MSC-based model for each focal species. Since each species has its own null distribution for correlations that is determined by the species tree (Fig. 3), each species has its own cutoff for significance of correlation coefficients. We include a table of all orthogroups that were significant at the *p <* 1*e −* 4 level, along with the top BLAST hit and putative gene functions (Tab. S1). Putative functions of these top hits were varied and included potentially relevant traits such as substrate specificity, growth, and hyphal extension.

Since our dataset includes only 8 focal species, we reasoned that we might have higher power and lower susceptibility to false positives if we pooled signals across species. While each focal species shares a single gene tree at each locus from which we calculate phylogenetic distances, competition indexes (RCC values) are measured independently for each pair of species, and hence only one measurement is shared between any pair of focal species across our estimates of correlation coefficients. Given this quasi-independence of the experimental data between focal species, we reasoned that genes that had a larger number of associations across focal species might be less likely to represent spurious associations. In our dataset, we observed one locus for which at least four of the eight focal species had corrected *p*-values under 1*e −* 2. In 100,000 independent coalescent simulations that used the same species tree and trait distribution, we never observed a locus for which four focal species had *p*-values under 0.01, suggesting that this observation is not likely under the null. We repeated this test for enrichment of loci harboring strong correlations across multiple species, using several different thresholds for significance and finding qualitatively similar results (Tab. S3).

The locus with the largest number of focal species with *p <* 1*e −* 4 is orthogroup 5719, with associations in both *Penicillium* SP. *mb* and *Penicillium polonicum*. It also had the most associations at the *α* = 5*e −* 2 level (in five out of eight of the species we studied; see Tab. S5). The top NCBI protein-protein BLAST hit for this protein was recoverin, which is a calcium signaling protein (Tamuli et al., 2011). In Fig. 5, we plot RCC values as a function of phylogenetic distance for each of the eight focal species, seven of which display a positive relationship between to phylogenetic distance and RCC. In contrast, the relationship between RCC and the species level phylogeny (based on naive linear models) was not significant for any single focal species at the *p* = 1*e −* 2 level (though *Penicillium verrucosum* narrowly missed this threshold with *p* = 0.0102; Fig. S4).

**Figure 5:**
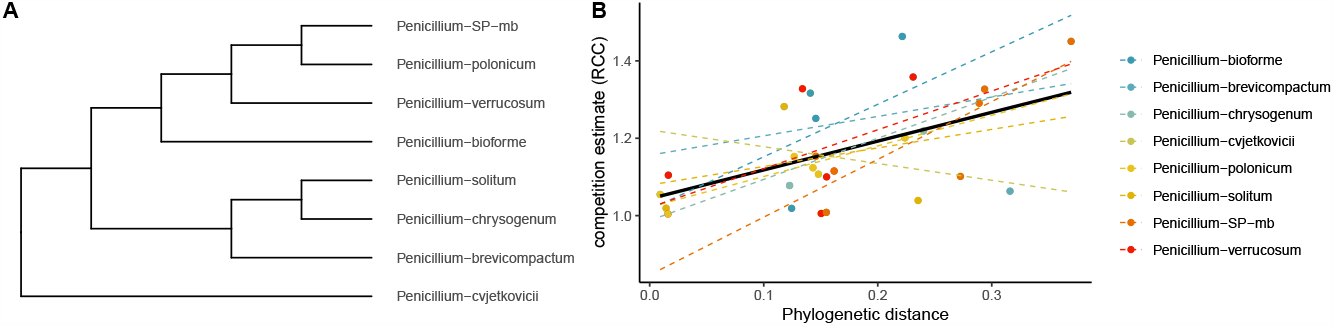
Gene tree and relationship between trait values and phylogenetic distance for the gene with the strongest phylogenomic signal. A: cladogram of the gene tree describing relationships between species for orthogroup 5719, for which the top `BLAST` hit was recoverin. B: Relationship between phylogenetic distance and RCC for each of the 8 species for this locus. Linear models were fit with `geom_smooth` in the `R` package `ggplot2`, using the method `lm`.

Since the number of associated genes exceeds the expectation under our null model, we reasoned that the set of associated genes might also be enriched for interesting biological functions. We used several different filtering strategies to create sets of associated genes and tested for pathway level enrichment in these sets using `FungiFun v2` (Priebe et al., 2015). We note that this analysis is somewhat limited from a statistical power perspective, because only a subset of genes could be mapped to the *P. chrysogenum* reference and only 1,892 genes have pathway-level annotations in `FungiFun v2`. While several pathways had low un-adjusted *p*-values, none survived an FDR-adjustment to account for multiple testing. Tab. S4 lists all pathways for which the FDR-adjusted *p*-value was below 0.25 and at least two genes in the pathway were in the foreground set. Interestingly, both DNA replication and phagosome-related genes were consistently among the top categories across several different strategies for determining the cutoff for significant associations between gene trees and RCC values. Upon closer inspection of the phagosome-related genes, we noted that several were subunits of the V-type proton ATPase (Pc13g08280, subunit C, chromosome I; Pc13g14670, V0, chromosome IV; Pc15g00440, subunit F, chromosome I; Pc21g12120, subunit G, location unknown; Pc22g24360, subunit E, chromosome I), and two others were subunits of Sec61 (Pc16g09660, subunit beta, chromosome I; Pc20g02810, subunit gamma, chromosome II). Defects in the function of V-type proton ATPases have been linked to growth defects in calcium-rich environments (Vasanthakumar and Rubinstein, 2020), while Sec61 has been implicated in calcium leakage in the endoplasmic reticulum (Lang et al., 2011). Although several of these genes fall on chromosome I, these patterns cannot be easily explained by genomic proximity because most of the gene pairs were megabases apart.

## Discussion

Relatedness between species might play a substantial role in determining community assembly dynamics and outcomes, but studies have found contrasting results when assessing the evidence for relatedness as a predictor of interspecific interaction strength. Many possible explanations for these discrepant findings have been proposed, including differences in phylogenetic scale (Graham et al., 2018), contrasting effects of relatedness on niche and fitness differences (Godoy et al., 2014), and the potential role of ecological filtering (Kivlin et al., 2014), to name a few. Here, we considered discordance between gene trees and species trees as a potential mechanism for obscuring the relationship between evolutionary history and competitive differences. Gene trees underlying genes that are causal for functional differences in competitive ability might be more strongly correlated with competition-related traits than the genome background. Our phylogenomic scan for such loci revealed several genes with strong correlations, in excess of what would be expected given a Multi-Species Coalescent-based null distribution. One gene, for which the top `BLAST` hit was recoverin, had branch lengths that were strongly correlated with competitive differences in five focal species, suggesting that it may be less likely to represent a spurious signal. Our results show that phylogenetic signal that would not be detected on the basis of species trees can be recovered using gene trees, and that our genomic scan for correlated gene trees can be used to infer loci that are associated with trait differences.

Association methods based on phylogenomic data have the potential to be used as tools to find the evolutionary and genomic underpinnings of interspecific trait variation. At their core, phylogenomic association methods operate under the assumption that a subset of genes in the genome are causal for trait differences across species, and that substitutions in these causal genes will be more informative about trait differences than the genome background. When this assumption is met, phylogenomic methods could come to different conclusions than prior approaches that examined only the species tree. In contrast, it will be more difficult to detect individual genes underlying traits that are highly polygenic with phylogenomic methods since each individual gene explains a small portion of the variance. In such cases, phylogenetic signal may often be easily detectable using the species tree even though the individual causal gene trees could be discordant, because the species tree represents a consensus over the gene trees. The extent to which discordance can explain discrepant results across prior studies therefore depends on both the polygenicity of traits and the extent of discordance in the investigated taxa. One potential step forward would be to re-examine prior studies to quantify the amount of discordance that would be expected given the species tree, and to consider whether studies with more expected discordance tend to find less evidence for phylogenetic signal. Even if discordance contributes to such contrasting patterns across studies, it does not preclude the possibility that other previously proposed explanations may also play a role.

Another potential step forward would be to apply phylogenomic association methods directly to trait data collected in prior studies of phylogenetic signal, when the relevant genomic sequence data is also available. In principle, our approach could be applied to any quantitative trait and taxa for which orthologs can be mapped. However, future studies employing such methods will need to carefully consider species sampling to maximize their effectiveness. Phylogenomic association approaches rely on signals of discordance and hence are most powerful when causal loci are discordant from the species tree. This suggests that species should be sampled such that internal branches of the tree are relatively short to maximize discordance, *i*.*e*. closely related groups of species should be sampled. Note that this also makes it easier to identify orthologs, since sequences are less diverged. As the number of species gets very large it becomes increasingly likely that in one or more species, any given ortholog will remain unmapped (or ambiguous), but it is straightforward to focus only on subsets of species at each locus for which an ortholog is known. It is also important that trait values have diverged enough across species such that there is substantial variation to detect – traits that are relatively invariant across the tree are less likely to result in positive associations. Ideally, trait values will have substantial variation both within and across the major clades of the species tree. Unknown aspects of the genomic basis for trait variation – such as polygenicity and the distribution of variance explained across genomic loci – will play a major role in determining the success or failure of such approaches (Fig. S2). If we hope to characterize the genomic basis of interspecific interactions or other ecologically relevant traits, it is important to choose traits that are closely related to the ecological processes of interest. In our study, the trait is derived from competition experiments on a cheese substrate that closely mimics the habitat of the eight *Penicillium* species. Controlled experiments in a shared environment are impossible for many species of interest. For example, it is currently infeasible to culture and experimentally manipulate many microbes, meaning that it would be difficult or impossible to obtain experimentally determined trait values such as those we used in this study. For these species, researchers can begin by leveraging existing phylogenomic datasets and prior research on traits that may be related to community composition and character displacement (Dayan and Simberloff, 2005).

When aspects of the environmental factors affecting growth or species interactions are known, it may be possible to relate the results of phylogenomic scans to these factors. For example, the gene with the most associations in our study is recoverin, a calcium signaling protein. Calcium signaling is crucial for regulating fungal growth, development, and maintenance of calcium homeostasis (Roy et al., 2021). Both substrate acidity (Wolfe et al., 2014) and the availability of soluble calcium (Amenu and Deeth, 2007) can change substantially as microbes grow during cheese production, possibly affecting the relative growth rates of fungal species growing on cheese substrates. Relating the results of phylogenomic scans back to known molecular functions of genes or environmental determinants of growth or reproduction could provide avenues for future hypotheses to be tested through experiments or observational data.

The most ideal endpoint of an association test is to experimentally validate (or falsify) the findings. While experimental validation is often challenging, tractable microbial systems can be particularly useful for testing hypotheses about community assembly (Cadotte et al., 2005; Vega and Gore, 2018; Chappell and Fukami, 2018). Phylogenomic scans (and association tests in general, including GWAS) cannot confirm a role for associated genes in the determination of trait values, but the results of these scans can be used as a screen to prioritize such outlier-loci for subsequent experiments to determine causality. For example, gene replacement studies can be used to remove and exchange orthologs of recoverin between species that have large differences in competitive abilities. Such experiments involve replacing a functional copy of a gene in one species with a sequence from another species, and comparing trait differences between species with and without the change in sequences. Such approaches can be easier to interpret than knockout studies, which can have substantial off-target fitness effects (for example, if a gene involved in competition is also essential for aspects of development or normal physiology). In the context of our eight *Penicillium* species, repeating the pairwise competition assay on mutant strains could confirm whether differences in competitive abilities are explained by sequence dissimilarity of recoverin. Gene-editing tools have become more widely available, allowing researchers to test evolutionary hypotheses in a much wider range of organisms (*e*.*g*., Karageorgi et al. 2019; Tannous et al. 2023).

Our phylogenomic scan is conceptually related to a recent approach known as `PhyloGWAS` (Pease et al., 2016), which scans the genome for genes that have a larger than expected number of nonsynonymouys substitutions that perfectly correlate with trait differences. `PhyloGWAS` detects associations between a set of nonsynonymous substitutions and a trait of interest, taking into account the background phylogeny by employing a multi-species coalescent (MSC) model. For each gene of interest, the number of nonsynonymous alleles that perfectly correlate with the trait is calculated, and a likelihood can be assigned to this substitution pattern by performing coalescent simulations given the species tree. This process naturally controls number of false-positive loci due to hemiplasy (*i*.*e*., patterns of discordance that can be well-explained by the null model of incomplete lineage sorting), and has been shown to be a powerful approach for detecting gene/trait associations. A minor limitation of `PhyloGWAS` led us to develop a new method for this study. As `PhyloGWAS` counts only perfectly correlated substitutions, it could be prone to decreased power in large sample sizes, where even one non-conforming lineage would discount the inclusion of a site of potential interest. Likewise, the inclusion of only perfectly correlated sites also limits the generalizability of the method to quantitative phenotypes, where there is no similar notion of a perfect correlation between trait values and substitution patterns. While these issues could be addressed by placing a lower threshold on the correlation-coefficient, there is not a clear *a priori* way to set such a threshold in general. Our approach of regressing gene tree branch lengths on trait values while controlling for the species tree under an MSC-based model extends this idea naturally to quantitative traits.

Like all correlative methods, our approach has several limitations. Although we took substantial care to infer an appropriate null distribution of correlation coefficients across gene trees, minor discrepancies between the null model and the true processes underlying variation in gene tree topology could cause mis-calibration and result in inflated test statistics that increase our false-positive rate. We note that our null is likely to be conservative, because we used summaries of the empirical distribution of correlation coefficients between gene trees and trait values to set the null. Any gene trees that have discrepant phylogenies due to processes that are not well explained by the multi-species coalescent, such as introgression events or misspecification of the true topology, will tend to increase the observed levels of discordance and hence make it more difficult to detect genes with aberrant topologies. In other words, our choice of null model should often result in a conservative reduction in power rather than an inflation in test statistics because the null is constructed from a gene set that includes both positive and negative genes, as well as introgressed genes. Like all genomic correlative methods, our approach comes with a substantial multiple testing burden. When we applied a strict Bonferroni cutoff, we retained only a handful of genes with exceptionally strong correlations (Tab. S1;Tab. S3). However, because the gene tree topologies in our dataset and others are unlikely to be completely independent (due to evolutionarily maintained syntenic blocks and linkage disequillibrium), a less stringent threshold could be applied by assessing the level of independence and the effective number of statistical tests. We also focused only on genes for which we could reliably infer a single-copy ortholog. Some genes of interest for competitive differences may have expanded in copy number and could have been excluded from our analyses.

Characterization and prediction of the mechanistic processes that determine community structure remains a substantial goal of community ecology, with much interest in how evolutionary processes may affect species interactions. Studies comparing trait differences to phylogenetic distances have provided evidence both for and against the hypothesis that competitive interactions are stronger between closely related species, with many potential resolutions proposed (*e*.*g*., Losos 1996; Godoy et al. 2014; Parmentier et al. 2014). How relatedness influences competition and coexistence between species likely depends on specific traits under selection, rather than aggregate trait differences. Genes encoding for specific traits influencing competition might not necessarily follow the same patterns of relatedness as the species tree. Our study thereby suggests a complementary role for discordance between gene trees and species trees in the interpretation of phylogenetic signal. We suggest that continued development of eco-evolutionary approaches that leverage the distinct evolutionary signatures in genes that are causal for traits, including traits that determine species interactions, will substantially enrich our understanding of the joint role of evolution and ecology in determining community assembly.

## Acknowledgements

This work was partially funded by a grant from the United States National Science Foundation (CAREER IOS/BIO 1942063) to B.E.W. The authors are grateful to Dr. Megan Biango-Daniels for collecting the original strains used in this experiment. We would also like to express our gratitude to Philip Carl Bentz for helpful discussions.

## Supporting Information

### Species sampling and selection

*Penicillium* strains were isolated on PCAMS (plate count agar with milk and salt; Cosetta and Wolfe 2020) from a cheese aging facility in the midwest region of the United States. Rind samples were stored in 1X phosphate buffered saline (PBS) for downstream processing. To culture fungi from samples, we diluted the cheese/slurry to 6-fold in PBS and plated out the 10^*−*3^ and 10^*−*4^ dilutions which we plated on PCAMS and supplemented with 50 mg/L chloramphenicol to inhibit bacterial growth. After maturation (*≈* 7 days), we harvested fungal spores which we were stored at -80 Celcius for molecular processing. Air samples were collected by capturing spores in the facility with an open petri dish containing cheese curd agar (CCA; Cosetta and Wolfe 2020), which were incubated at room temperature and subsequently processed like the cheese rind samples. For *in vitro* competition experiments, we chose a subset of 8 species which are phylogenetically and morphologically diverse (Bodinaku et al., 2019). The eight species are *Penicillium cvjetkovicii, Penicillium breivcompactum, Penicillium* SP. *mb, Penicillium bioforme, Penicillium polonicum, Penicillium solitum, Penicillium verrucosum*, and *Penicillium chrysogenum*.

### Competition experiments

To construct 28 unique interspecific pairwise microcosms representing all pairs of the eight focal species in our study, we standardised cell concentrations to 20 colony forming units (CFUs)/μL following methods as described earlier (Cosetta and Wolfe, 2020). Next, we spot-plated 10μL of standardized innoculum for competing strains 2.5 cm from opposing edges of petri dishes containing CCA. For each strain, we included a pair of intraspecific competition as a control, with the competitor replaced by another replicate of the same strain at the same cell density. All microcosms were repeated for three biological replicates. Plates were incubated in low light levels at 14 Celcius — simulating conditions of a cheese-aging facility — for 21 days. After incubation, all microcosms were scanned using an Epson Perfection V37 document scanner to acquire standardized images of competitive outcomes.

To quantify competitive outcomes, we uploaded scanned images to `ImageJ` (Abràmoff et al., 2004) and used the “Polygon” function to measure the area occupancy of each strain per microcosm. Using the area occupancy data, we calculated a competitive index for each pairwise interaction. We used the Relative Crowding Coefficient (RCC) as a measure of competitive ability, which was originally developed for understanding competitive interactions between target crops and weeds (De Wit, 1960). RCC is calculated as 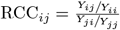 of species *i* when grown with species *j*, where *Y* is the area of occupancy after growing together for a fixed period of time. *Y*_*ji*_ corresponds to the area occupied by species *j* when grown with competitor species *i*. All measurements in this study were taken after a period of 21 days. Since input cell densities between competitors were approximately equal, an RCC of greater than 1 indicates that species *i* is more competitive than *j*, whereas with an RCC score lower than 1, *j* is more competitive than *i*. We performed three replicate experiments for each pair of competitors, and took the average value of RCC across the replicates as our measure of competitive ability.

### Genomic sequencing

DNA was extracted using the Powersoil Pro DNA extraction kit (MoBio, Carlsbad, CA, USA), following kit instructions. Library preparation was performed with NEBNext^®^ Ultra™ II DNA Library Prep (New England Biolabs, Ipswich, MA, USA), which were sequenced at Tufts University Core Facility Genomics on an Illumina NextSeq550. Low quality reads were trimmed and removed for *de novo* assembly using `GenSAS` (Humann et al., 2019), while `Augustus` (Stanke et al., 2004) was use to annotate coding regions in these fungal genomes.

### Phylogenomic analyses

To identify orthologous genes between fungal strains, we used `OrthoFinder` version 2.5.4 (Emms and Kelly, 2019), with default settings. We generated gene trees of orthologous genes using `FastTree` version 2.1.10 (Price et al., 2010), and performed a multiple sequence alignment on subsequent gene trees using `MAFFT` version 7.481 (Katoh and Standley, 2013). The alignments were concatenated together and `RAxML-NG` version 1.1.0 was used to infer the species tree using model `LG+G8+F`, which uses a discrete Gamma with eight categories and the evolutionary model proposed by Le and Gascuel (Le and Gascuel, 2008). Only single-copy orthologous genes, for which we identified a single ortholog in all 8 fungal species, were included in our analyses, of which there were 4,544. Gene trees and sequences for all of these genes are available at github.com/uricchio/ILSComp.

### Simulations

We simulated quantitative traits under a simple evolutionary model of trait differentiation using the `Python` package `DendroPy` (Sukumaran and Holder, 2010). Given the phylogeny of the species, we first simulate a gene tree for a causal gene under the multi-species coalescent model. We then simulate a quantitative trait evolving neutrally on the causal gene tree through the accumulation of substitutions. We fixed the species tree topology in our simulations to the inferred topology for the eight *Penicillium* species in our study.

For our quantitative trait model, we suppose that the number of substitutions occurring on a branch of length *T* is Poisson distributed with parameter *Tθ*, where *θ* is the number of substitutions expected per unit of time *T* . We suppose that each substitution in a causal gene has an additive effect on the trait of interest, with effect sizes that are drawn from a standard normal distribution.

We compared three broad scenarios in our simulations, which correspond to high, moderate, and low discordance between gene trees and species trees. In these simulations, we suppose that each branch of the species tree has the same population size. We set the population size parameter to be very low relative to branch lengths such that most gene trees have the same topology as the species tree for low discordance scenarios, while population size is very large relative to branch lengths for high discordance scenarios. Parameters for these simulations are given in Tab. S2. Note that since the gene trees and species trees we use in the text have branch lengths scaled in units of substitutions per base-pair, the population sizes used in the simulations do not translate to numbers of individuals, but rather the rate of coalescence relative to the branch lengths.

In general, analyses of statistical power and Type-I error rates under our model are both dependent on the model parameters for the quantitative trait evolution and the topology of the species tree. Our trait model does not account for selection, interactions between loci, unknown distributions of mutational effects, and other complicating factors. As such, we take these simulations only as proof of concept and not a full exploration of the statistical performance of our approach under plausible evolutionary models. The analyses herein cannot be taken as general statements about power, but rather comparisons across a range of scenarios that encapsulate high and low discordance between gene trees and species trees. Software implementing these analyses is available at github.com/uricchio/ILSComp.

### Statistical inference of phylogenetic signal

We compared three inference approaches in this study. First, we performed naive correlation analyses between gene tree branch lengths and squared differences in trait values, without correcting for the structure of the species tree. Second, we used Phylogenetic Least Squares (PGLS) as implemented in `ape` version 5.6-2 and `geiger` version 2.10.0 (Paradis and Schliep, 2019; Pennell et al., 2014) to regress squared trait differences on distances calculated with gene trees, taking the species tree as a co-variate. Third, we used coalescent simulations to generate a null distribution over the correlation between trait differences and gene tree branch lengths, given a species tree and a set of trait values at the leaves of the tree. We assumed a Brownian model of trait evolution in our PGLS analyses. We assessed the ability of the latter two approaches to correct for the expected correlation in trait values due to the shared phylogeny and identify significant outliers. We describe the third approach (MSC-based simulations) in more detail in the next section.

We calculated phylogenetic distances between species in both real and simulated gene trees using `DendroPy`. When we regress trait differences on phylogenetic distances in simulated data, we compare the squared difference between trait values to the phylogenetic distance because the expectation of the signed difference in trait values is zero under a neutral evolutionary model. When we consider real *Penicillium* gene trees and traits, we use an experimentally determined measure of competition strength as the trait values (RCC, see Competition Experiments below) for each species pair as the trait value. Because RCC for a pair of species *i* and *j* has the property that RCC_*i←j*_ = ^1^*/*RCC_*j←i*_, we use whichever of the RCC values is greater than one in our analyses. This is analagous to using the squared trait differences as we do in the simulations, since RCC = 1 indicates competitive equivalence and larger RCC values suggest larger overall differences in competitive ability. In other words, we are not concerned with the identity of the better competitor, but rather the magnitude of competitive differences between species as a function of phylogenetic distance. We used the function `linregress` in the `Python` package `scipy.stats` to calculate correlation coefficients and uncorrected *p*-values for the association between trait values and phylogenetic distances.

In all regression analyses performed in this study, we separately regressed distances between a single focal species and each of its seven competitors against the competition index (RCC) for each competitor with the focal species, rather than combining all eight species into a single analysis with all 28 pairwise measures included. We do not include self comparisons in any of our regressions, because they always have an evolutionary/trait distance of 0 by definition which could bias analyses. We reasoned that some genes might be of importance in determining only a subset of competitive outcomes, and that combining all 28 pairs could therefore decrease our power to detect some associations. This does come at some cost, in the sense that we could lose power to detect genes with weak shared effects across all (or most) of the pairs in our study. To address this limitation, we also performed tests of enrichment in which we assess the evidence for a larger than expected number of genes with multiple associations across focal species using a hypergeometric model and coalescent simulations (see Tab. 1 and Tab. S3). Note that our approach could easily be used for applications in which it is desirable to analyze all species pairs simultaneously with only minor modifications to the software.

### Determining a null distribution of *ρ* under the MSC

To calculate the null distribution of correlation coefficients between neutrally evolving gene trees and trait values under the coalescent model, we simulate neutral gene trees on the species tree given a fixed distribution of trait values. These fixed trait values correspond either to the simulated trait values for analyses of statistical performance, or to the experimentally determined RCC values for each pair of species. We then determine a null distribution of correlation coefficients between simulated phylogenetic distances and the observed trait values by regressing the simulated non-causal branch lengths of the neutral gene trees against the trait values.

While it is straightforward to perform these simulations when all relevant model parameters are known, we do not have empirically estimated population sizes for each species of *Penicillium* in our study. Since population sizes determine the rate of coalescence along a branch of the phylogenetic tree, we need this parameter value to perform our simulations and estimate the null distribution of correlation coefficients. As the amount of discordance between gene trees and species trees generally increases as a function of population size, we can use an estimate of the discordance in a set of gene trees to determine an appropriate population size for our simulations. To determine an appropriate population size, we simulate sets of gene trees at a range of different population sizes and calculate the mean and standard deviation of correlation coefficients between the branch lengths and trait values for the simulated gene trees. We then minimize the Euclidean distance between these simulated summary statistics and the same values in our observed dataset. We take the population size that minimizes the difference between the observed and simulated summary statistics as an estimate of the population size (Fig. S3).

We used our simulated null distributions of correlation coefficients to determine *p*-values of correlation coefficients. We use one-sided *p*-values in this study because we are only interested in the subset of loci where correlations between gene tree branches and traits are greater than expected. The *p*-values were determined by computing the fraction of simulated gene trees that had correlation coefficients in excess of the observed correlation coefficient for a given gene tree in our empirical dataset.

### Simulations of traits with polygenic architecture

We assessed the performance of our approach for traits with polygenic architecture (*i*.*e*., more than one locus harbors substitutions that affect trait values). To perform this analysis, we simulated species trees with various numbers of lineages under a birth-death process using `DendroPy`. We then simulated causal gene trees on the simulated species trees in a manner exactly analogous to the high discordance simulations we performed for monogenic traits. The underlying evolutionary model was identical to the model described in the simulation methods im the main text, but trait values on the tips of the tree are determined by summing the effects of each individual causal gene. We supposed that discordance was very high for these simulations. We then calculated the power of our method to detect associations amongst the set of genes. Code implementing the simulations is available at https://uricchio.github.io/ILSComp. Results of these simulations are plotted in Fig. S2.

### Tests of functional enrichment

We performed several tests for pathway-level enrichment among sets of genes that were positively correlated with RCC using KEGG annotations. To perform these tests, we used the `FungiFun 2.2.8` (available as a webtool at https://elbe.hki-jena.de/fungifun/; Priebe et al. 2015). We downloaded 12,789 reference *P. chrysogenum* amino acid sequences from `UniProt`, taxon ID 500485 (https://www.uniprot.org/taxonomy/500485). We then used `OrthoFinder` to map orthologs between our sequenced *P. chrysogenum* isolate and this *P. chrysogenum* reference. We used the set of one-to-one orthologs between these datasets to create a “background” gene set, *i*.*e*. a set of all genes that could possibly be considered in our test for functional enrichment. Of the 4,544 single copy orthologs, 3,629 were mapped uniquely to genes in the *P. chrysogenum* reference genome to form this background set. We created several test sets of “foreground” genes that corresponded to different strategies for detecting associations (*i*.*e*. different *p*-value thresholds and number of associated species), as described in Tab. S4. We note that these tests of enrichment should be taken with caution for multiple reasons, perhaps most importantly because functionally related genes may be located proximally to each other on the genome, meaning that their gene trees may not be fully independent. This could violate the assumptions of the hypergeometric distribution, which was used to calculate the statistical significance of pathway enrichment. When possible, we used the aforementioned *P. chrysogenum* `UniProt` reference sequence and the `EnsemblFungi` genome browser (http://fungi.ensembl.org/Penicillium_chrysogenum_gca_000710275/) to determine the chromosomal location of each associated gene.

**Figure S1:**
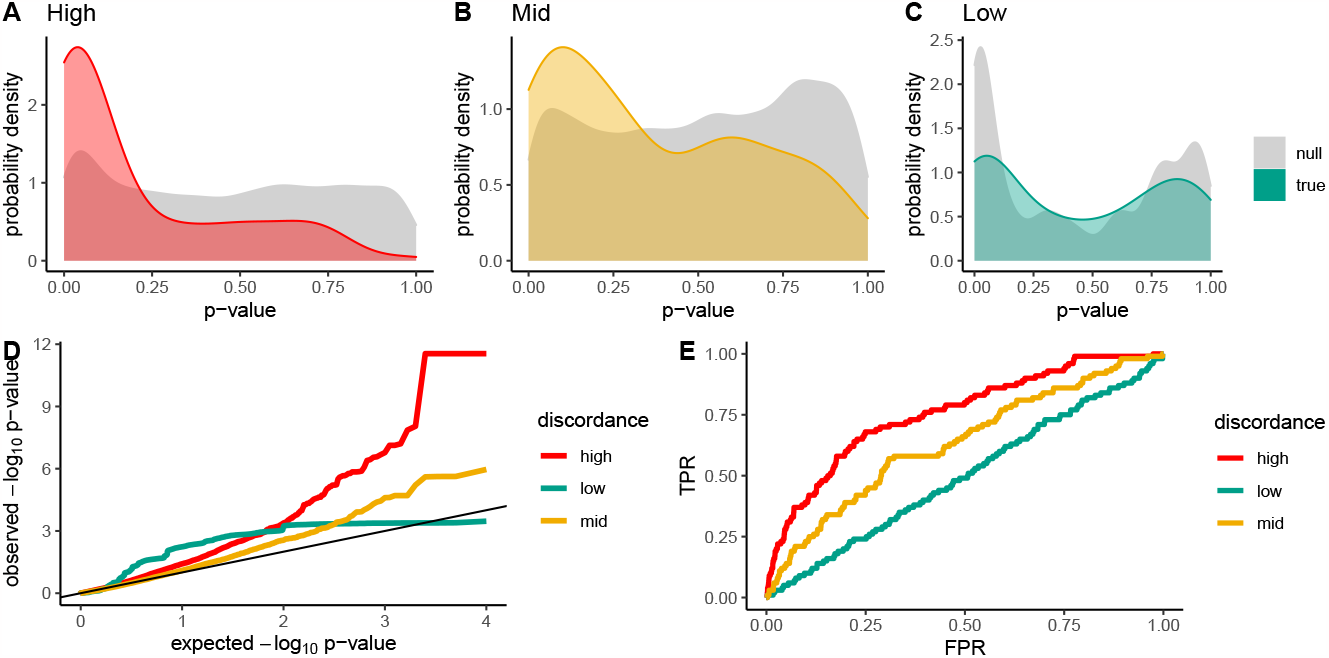
A-C: Probability density of *p*-values obtained for the correlation between phylogenetic distances and simulated trait differences in trait values under the “null” (*i*.*e*., random gene trees) and for the “true” causal gene tree for each of three levels of discordance, when PGLS is used to perform the regression. Each panel in this figure corresponds only to differences between *P. bioforme* and each of its competitor species because each species has it’s own null distribution. D: QQ-plot for *p*-values obtained under the null for high, low, and moderate gene/species discordance. E: ROC curve for identification of causal genes based on *p*-values of the correlation between gene tree branch lengths and pairwise trait differences

**Figure S2:**
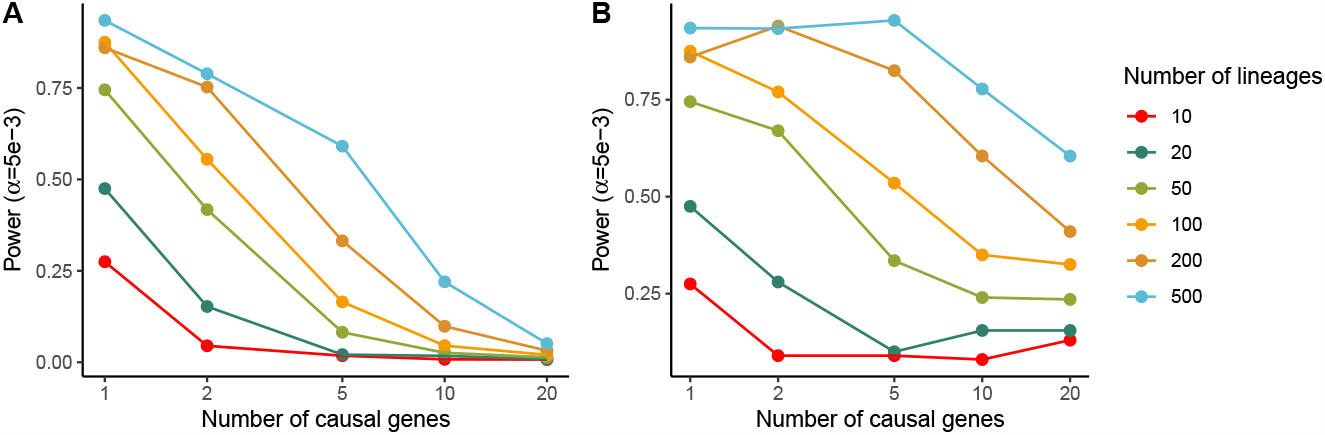
A: Power at the *α* = 5*e −*3 level to detect causal genes for quantitative trait as a function of the number of species and the number of causal genes. Power generally increases as the number of species increases and declines as the number of causal genes increase. Note that these simulations suppose high levels of discordance between gene trees and the species tree (see Supporting Information). B: Power to detect at least one causal gene out of a set of causal genes.

**Figure S3:**
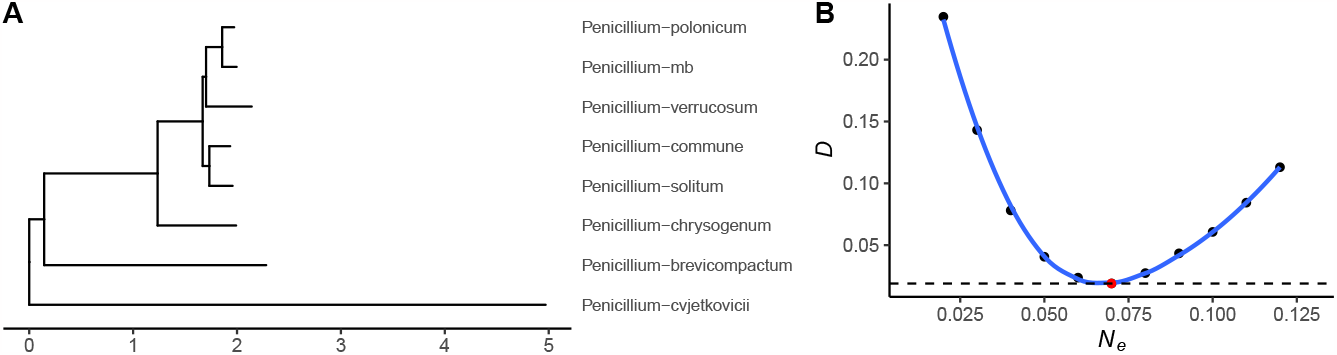
A: Species tree scaled by coalescent units. B. Inference of the population size used to scale the tree in A, which is a critical parameter for determining a null distribution over correlation coefficients. We minimized *D* as a function of *N*_*e*_, where *D* the Euclidean distance between the mean and standard deviation of the simulated distribution of correlation coefficients and the observed distribution of correlation coefficients (*i*.*e, D* = ∑_*i*_ (*μ*_*o−*_*μ*_*s*_)^2^ + (*σ*_*o−*_*σ*_*s*_)^2^, where *μ* is the mean and *σ* is the standard deviation, and the subscript *o* indicates observed values and *s* indicates simulated values and *i* indexes over species included in the analysis). For each value of *N*_*e*_ we simulate gene trees given the species tree, calculate *D* for the same trait distribution observed in our study, and then minimize *D* to find the best-fitting value of *N*_*e*_.

**Figure S4:**
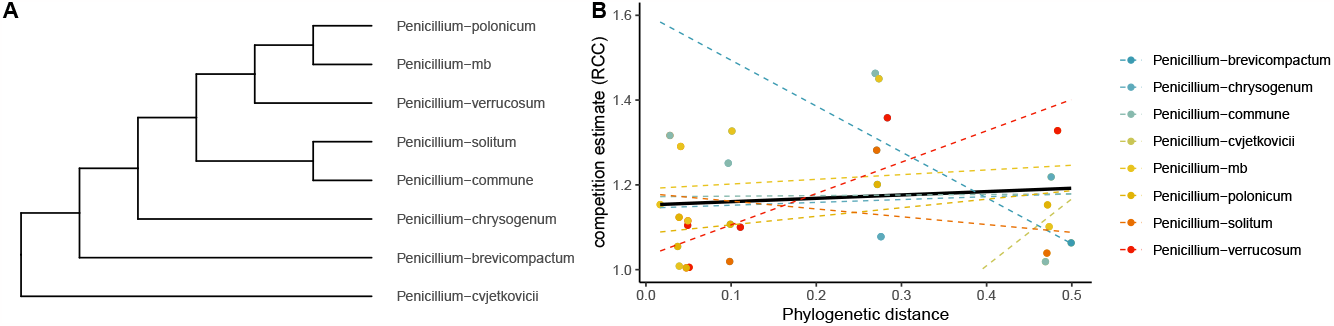
Relationship between phylogenetic distance and RCC for each of the 8 species for the species tree. Linear models were fit with `geom_smooth` in the `R` package `ggplot2`, using the method `lm`.

**Table S1:**
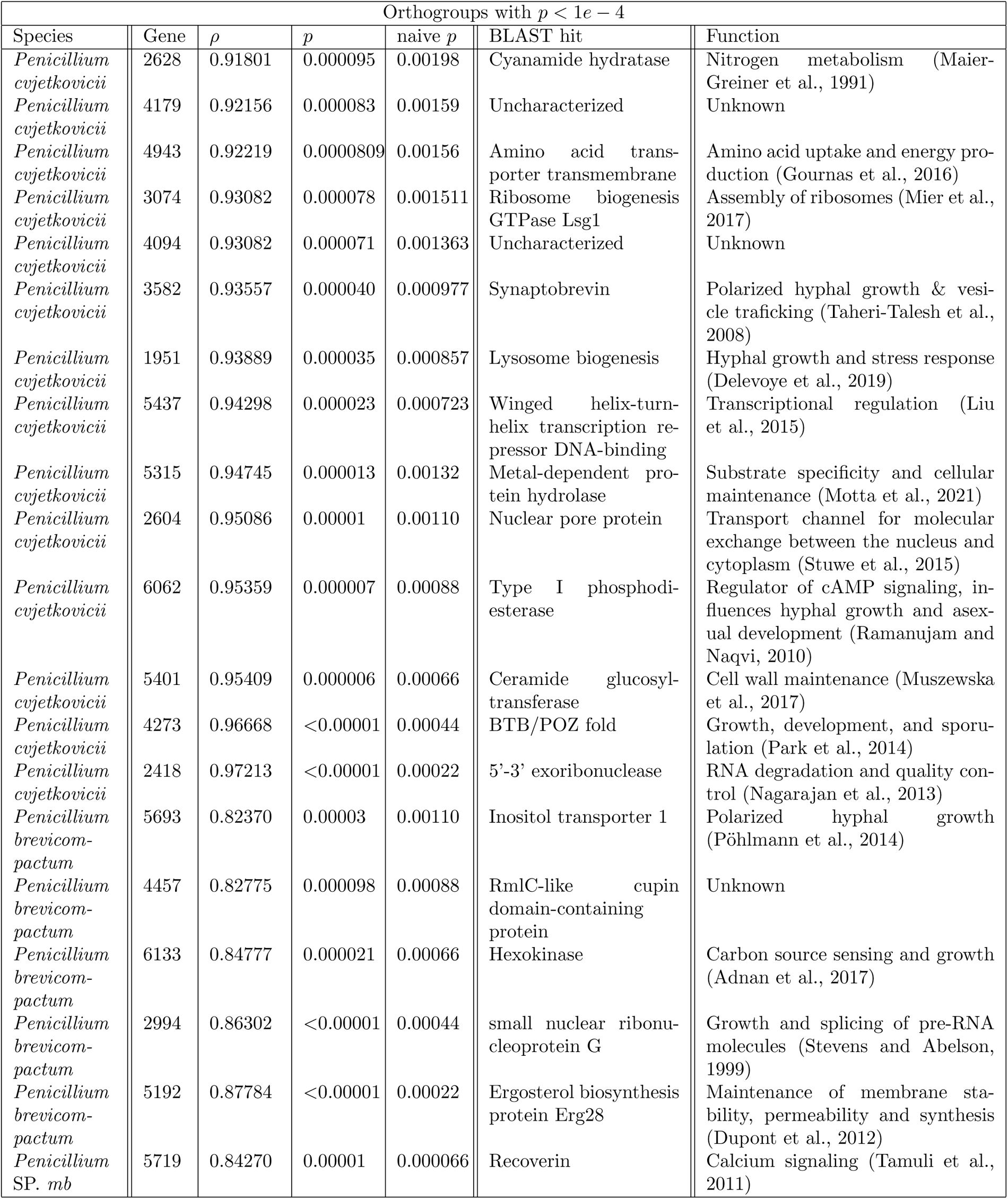
Othogroups with *p <* 1*e −*4. Naive *p* refers to the *p*-value given by a naive linear model, while *p* corresponds to our estimate of the probability of the observed correlation under the coalescent model. The Gene column indicates the number of the orthogroup in our dataset, which can be accessed at github.com/uricchio/ILSComp.

| Orthogroups with $p < 1e - 4$ | | | | | | |
| --- | --- | --- | --- | --- | --- | --- |
| Species | Gene | $\rho$ | $p$ | naive $p$ | BLAST hit | Function |
| *Penicillium cvjetkovicii* | 2628 | 0.91801 | 0.000095 | 0.00198 | Cyanamide hydratase | Nitrogen metabolism (Maier-Greiner et al., 1991) |
| *Penicillium cvjetkovicii* | 4179 | 0.92156 | 0.000083 | 0.00159 | Uncharacterized | Unknown |
| *Penicillium cvjetkovicii* | 4943 | 0.92219 | 0.0000809 | 0.00156 | Amino acid transporter transmembrane | Amino acid uptake and energy production (Gournas et al., 2016) |
| *Penicillium cvjetkovicii* | 3074 | 0.93082 | 0.000078 | 0.001511 | Ribosome biogenesis GTPase Lsg1 | Assembly of ribosomes (Mier et al., 2017) |
| *Penicillium cvjetkovicii* | 4094 | 0.93082 | 0.000071 | 0.001363 | Uncharacterized | Unknown |
| *Penicillium cvjetkovicii* | 3582 | 0.93557 | 0.000040 | 0.000977 | Synaptobrevin | Polarized hyphal growth & vesicle trafficking (Taheri-Talesh et al., 2008) |
| *Penicillium cvjetkovicii* | 1951 | 0.93889 | 0.000035 | 0.000857 | Lysosome biogenesis | Hyphal growth and stress response (Delevoye et al., 2019) |
| *Penicillium cvjetkovicii* | 5437 | 0.94298 | 0.000023 | 0.000723 | Winged helix-turn-helix transcription repressor DNA-binding | Transcriptional regulation (Liu et al., 2015) |
| *Penicillium cvjetkovicii* | 5315 | 0.94745 | 0.000013 | 0.00132 | Metal-dependent protein hydrolase | Substrate specificity and cellular maintenance (Motta et al., 2021) |
| *Penicillium cvjetkovicii* | 2604 | 0.95086 | 0.00001 | 0.00110 | Nuclear pore protein | Transport channel for molecular exchange between the nucleus and cytoplasm (Stuwe et al., 2015) |
| *Penicillium cvjetkovicii* | 6062 | 0.95359 | 0.000007 | 0.00088 | Type I phosphodiesterase | Regulator of cAMP signaling, influences hyphal growth and asexual development (Ramanujam and Naqvi, 2010) |
| *Penicillium cvjetkovicii* | 5401 | 0.95409 | 0.000006 | 0.00066 | Ceramide glucosyltransferase | Cell wall maintenance (Muszewska et al., 2017) |
| *Penicillium cvjetkovicii* | 4273 | 0.96668 | <0.00001 | 0.00044 | BTB/POZ fold | Growth, development, and sporulation (Park et al., 2014) |
| *Penicillium cvjetkovicii* | 2418 | 0.97213 | <0.00001 | 0.00022 | 5'-3' exoribonuclease | RNA degradation and quality control (Nagarajan et al., 2013) |
| *Penicillium brevicompactum* | 5693 | 0.82370 | 0.00003 | 0.00110 | Inositol transporter 1 | Polarized hyphal growth (Pöhlmann et al., 2014) |
| *Penicillium brevicompactum* | 4457 | 0.82775 | 0.000098 | 0.00088 | RmlC-like cupin domain-containing protein | Unknown |
| *Penicillium brevicompactum* | 6133 | 0.84777 | 0.000021 | 0.00066 | Hexokinase | Carbon source sensing and growth (Adnan et al., 2017) |
| *Penicillium brevicompactum* | 2994 | 0.86302 | <0.00001 | 0.00044 | small nuclear ribonucleoprotein G | Growth and splicing of pre-RNA molecules (Stevens and Abelson, 1999) |
| *Penicillium brevicompactum* | 5192 | 0.87784 | <0.00001 | 0.00022 | Ergosterol biosynthesis protein Erg28 | Maintenance of membrane stability, permeability and synthesis (Dupont et al., 2012) |
| *Penicillium SP. mb* | 5719 | 0.84270 | 0.00001 | 0.000066 | Recoverin | Calcium signaling (Tamuli et al., 2011) |

**Table S2:** Parameters relevant to our simulations of gene-tree, species-tree discordance. Lower population sizes (relative to branch lengths) result in increased levels of discordance. *θ* = 2*NuL* corresponds to the mutation rate over a gene of length *L*, such that the expected number of substitutions along a simulated branch of length *T* is *Tθ*. Note that the branch lengths are scaled in units of substitutions per site, so the population sizes here do not correspond to numbers of individuals. We allowed the mutation rate to decrease with population size, because larger population sizes result in longer branches for gene trees, and hence would have more substitutions at a given value of *θ*.

| Simulation parameters |  |  |
| --- | --- | --- |
| Model | $\theta = 2Nu$ | Population size ($N$) |
| Low discordance | $10^4$ | $10^{-3}$ |
| Mid discordance | $10^2$ | $10^{-1}$ |
| High discordance | $10^{-4}$ | $10^5$ |

**Table S3:** Results of enrichment tests with various parameters. “Sig. threshold” corresponds to the threshold *p*-value below which a single gene:trait association was considered to be significant, and “No. of focal species” is the number of species for which a locus needed to have *p*-values below the threshold to be considered a significant hit. The Null column indicates the number of simulations (out of 100,000) for which we observed a positive association given the parameters, and Observed indicates the number of associations we observed (out of 4,544) in our real dataset. The *p* column indicates the probability of the observed data given the simulations under a hypergeometric model (function `dhyper` in `R`). The probability of 1 observation out of 4,544 given a background of 0 out of 100,000 is not well defined for the hypergeometric (*i*.*e*., the values in the first row), so we conservatively estimated this probability supposing that one observation was expected in 100,000 trials resulting in *p <* 0.045.

| Tests for enrichment in the number of associated genes |  |  |  |  |
| --- | --- | --- | --- | --- |
| Sig. threshold | No. of focal species | Null | Observed | $p$-value |
| 0.01 | 4 | 0 | 1 | <0.045 |
| 0.01 | 3 | 91 | 6 | 0.113 |
| 0.05 | 4 | 812 | 21 | 0.007 |
| 0.05 | 3 | 3,450 | 151 | 0.03 |

**Table S4:** Results of tests for KEGG pathway enrichment. Each row represents a pathway. The values in the first two columns were used to determine the set of “foreground” genes for the enrichment test. We list only pathways for which the FDR-adjusted p-value was below 0.25 and at least two genes in the pathway were in the foreground set. “Single species sig. threshold” corresponds to the threshold *p*-value for associations (*i*.*e*., a single gene-species pair was considered significant if its p-value was below the threshold), and “Focal species” is the number of species that needed to have *p*-values below the threshold at a single locus in order for the locus to be included in the foreground set of genes. “Pathway” is the name of the putatively enriched pathway, and “Genes” is the set of genes that were among the foreground set. The un-adjusted *p*-value column indicates the probability of the observed pathway enrichment under a hypergeometric model, while the FDR adjustment represents a correction for multiple testing. These analyses were performed with `FungiFun v.2` (Priebe et al., 2015), and the gene identifiers correspond to the *P. chrysogenum* reference identifiers in the `FungiFun v.2` database.

| Tests of functional enrichment |  |  |  |  |  |
| --- | --- | --- | --- | --- | --- |
| Single species sig. threshold | Focal species | Pathway | Genes | Un-adjusted p-value | FDR-adjusted p-value |
| 0.0005 | 1 | Phagosome | Pc15g00440, Pc20g02810 | 0.0254 | 0.22903 |
| 0.005 | 1 | Phagosome | Pc13g08280, Pc13g14670, Pc15g00440, Pc16g09660, Pc20g02810, Pc21g12120 | 0.00261 | 0.1225 |
| 0.01 | 2 | Phagosome | Pc13g08280, Pc13g14670, Pc15g00440, Pc20g02810 | 0.004733925 | 0.1420177 |
| 0.05 | 2 | DNA replication | Pc12g04440, Pc13g07420, Pc16g10450, Pc20g14580, Pc21g08060, Pc21g10710, Pc21g11440, Pc21g14450, Pc21g15650, Pc21g21700 | 0.001999 | 0.1339 |
| 0.05 | 2 | Phagosome | Pc13g08280, Pc13g14670, Pc15g00440, Pc16g09660, Pc18g01990, Pc20g02810, Pc21g12120, Pc22g24360 | 0.007726 | 0.242686 |
| 0.05 | 2 | Spliceosome | Pc12g04780, Pc13g03680, Pc13g04070, Pc13g06570, Pc13g10450, Pc13g11030, Pc16g04930, Pc16g04940, Pc20g01340, Pc20g09280, Pc20g10480, Pc20g15290, Pc21g00700, Pc21g04460, Pc22g04980 | 0.01087 | 0.242686 |
| 0.05 | 3 | DNA replication | Pc12g04440, Pc13g07420, Pc16g10450, Pc20g14580, Pc21g10710, Pc21g11440, Pc21g15650, Pc21g21700 | 0.00063943 | 0.0358 |
| 0.05 | 3 | Phagosome | Pc13g08280, Pc13g14670, Pc16g09660, Pc18g01990, Pc20g02810, Pc21g12120 | 0.005833 | 0.1633 |
| 0.05 | 3 | Mismatch repair | Pc12g04440, Pc13g07420, Pc20g14580, Pc21g11440, Pc21g21700 | 0.01304 | 0.2436 |
| 0.05 | 4 | Mismatch repair | Pc20g14580, Pc21g21700 | 0.0186 | 0.16729 |
| 0.05 | 4 | Spliceosome | Pc16g04930, Pc20g09280, Pc22g04980 | 0.0252 | 0.16729 |
| 0.05 | 4 | DNA replication | Pc20g14580, Pc21g21700 | 0.0358 | 0.16729 |

**Table S5:** Correlation coefficients and p-values for each focal species. *ρ*_*s*_ and *p*_*s*_ correspond to the correlation coefficient and *p*-value for the linear model *RCC D*_*s*_, where *RCC* is the competition coefficient as defined in the text for the focal species as compared to a competitor, and *D*_*s*_ is the evolutionary distance given the species tree. *ρ*_*R*_ is the correlation coefficient for the linear model *RCC D*_*R*_, where *RCC* is the competition coefficient as defined in the text for the focal species as compared to a competitor, and *D*_*R*_ is the evolutionary distance given the gene tree for the recoverin locus. *p*_*R*_ corresponds to the corrected *p*-value that we obtain by comparing the observed correlation coefficient to a multi-species coalescent null model. Data corresponding to these linear regressions are plotted in Fig. S4 (species tree) and Fig. 5 (recoverin gene tree).

| Correlation coefficients and p-values |  |  |  |  |
| --- | --- | --- | --- | --- |
| Species | $\rho_s$ | $\rho_R$ | $p_s$ | $p_R$ |
| *Penicillium cvjetkovicii* | 0.223 | -0.306 | 0.6302 | 0.945406 |
| *Penicillium brevicompactum* | -0.553 | 0.281 | 0.1971 | 0.057024 |
| *Penicillium bioforme* | 0.0083 | 0.602 | 0.986 | 0.003599 |
| *Penicillium* SP. *mb* | 0.119 | 0.843 | 0.799 | 0.000066 |
| *Penicillium chrysogenum* | 0.8417 | 0.857 | 0.09368 | 0.000391 |
| *Penicillium polonicum* | 0.515 | 0.922 | 0.2368 | 0.000147 |
| *Penicillium verrucosum* | 0.871 | 0.557 | 0.0107 | 0.864137 |
| *Penicillium solitum* | -0.227 | 0.302 | 0.6245 | 0.019773 |

## References

M. D. Abràmoff, P. J. Magalhães, and S. J. Ram. Image processing with ImageJ. Biophotonics International, 11(7): 36–42, 2004.

M. Adnan, W. Zheng, W. Islam, M. Arif, Y. S. Abubakar, Z. Wang, and G. Lu. Carbon catabolite repression in filamentous fungi. International Journal of Molecular Sciences, 19(1):48, 2017.

B. Amenu and H. Deeth. The impact of milk composition on cheddar cheese manufacture. Australian Journal of Dairy Technology, 62(3):171, 2007.

I. Bodinaku, J. Shaffer, A. B. Connors, J. L. Steenwyk, M. N. Biango-Daniels, E. K. Kastman, A. Rokas, A. Robbat, and B. E. Wolfe. Rapid phenotypic and metabolomic domestication of wild penicillium molds on cheese. mBio, 10 (5):e02445–19, 2019.

Å. Brännström and D. J. Sumpter. The role of competition and clustering in population dynamics. Proceedings of the Royal Society B: Biological Sciences, 272(1576):2065–2072, 2005.

M. W. Cadotte, J. A. Drake, and T. Fukami. Constructing nature: laboratory models as necessary tools for investigating complex ecological communities. Advances in Ecological Research, 37:333–353, 2005.

J. F. Cahill Jr, S. W. Kembel, E. G. Lamb, and P. A. Keddy. Does phylogenetic relatedness influence the strength of competition among vascular plants? Perspectives in Plant Ecology, Evolution and Systematics, 10(1):41–50, 2008.

J. Cavender-Bares, K. H. Kozak, P. V. Fine, and S. W. Kembel. The merging of community ecology and phylogenetic biology. Ecology Letters, 12(7):693–715, 2009.

C. R. Chappell and T. Fukami. Nectar yeasts: a natural microcosm for ecology. Yeast, 35(6):417–423, 2018.

C. M. Cosetta and B. E. Wolfe. Deconstructing and reconstructing cheese rind microbiomes for experiments in microbial ecology and evolution. Current Protocols in Microbiology, 56(1):e95, 2020.

T. J. Davies, N. Cooper, J. A. F. Diniz-Filho, G. H. Thomas, and S. Meiri. Using phylogenetic trees to test for character displacement: a model and an example from a desert mammal community. Ecology, 93(p8):S44–S51, 2012.

T. Dayan and D. Simberloff. Ecological and community-wide character displacement: the next generation. Ecology letters, 8(8):875–894, 2005.

C. T. De Wit. On competition. Pudoc, Wageninoen, The Netherlands, 1960.

J. H. Degnan and N. A. Rosenberg. Gene tree discordance, phylogenetic inference and the multispecies coalescent. Trends in Ecology & Evolution, 24(6):332–340, 2009.

C. Delevoye, M. S. Marks, and G. Raposo. Lysosome-related organelles as functional adaptations of the endolysosomal system. Current opinion in cell biology, 59:147–158, 2019.

S. Dupont, G. Lemetais, T. Ferreira, P. Cayot, P. Gervais, and L. Beney. Ergosterol biosynthesis: a fungal pathway for life on land? Evolution, 66(9):2961–2968, 2012.

I. Ebersberger, P. Galgoczy, S. Taudien, S. Taenzer, M. Platzer, and A. Von Haeseler. Mapping human genetic ancestry. Molecular Biology and Evolution, 24(10):2266–2276, 2007.

N. B. Edelman and J. Mallet. Prevalence and adaptive impact of introgression. Annual Review of Genetics, 55: 265–283, 2021.

S. V. Edwards, Z. Xi, A. Janke, B. C. Faircloth, J. E. McCormack, T. C. Glenn, B. Zhong, S. Wu, E. M. Lemmon, A. R. Lemmon, et al. Implementing and testing the multispecies coalescent model: a valuable paradigm for phylogenomics. Molecular Phylogenetics and Evolution, 94:447–462, 2016.

D. M. Emms and S. Kelly. OrthoFinder: phylogenetic orthology inference for comparative genomics. Genome Biology, 20:1–14, 2019.

J. Felsenstein. Phylogenies and the comparative method. The American Naturalist, 125(1):1–15, 1985.

N. Galtier and V. Daubin. Dealing with incongruence in phylogenomic analyses. Philosophical Transactions of the Royal Society B: Biological Sciences, 363(1512):4023–4029, 2008.

O. Godoy, N. J. Kraft, and J. M. Levine. Phylogenetic relatedness and the determinants of competitive outcomes. Ecology letters, 17(7):836–844, 2014.

C. Gournas, M. Prévost, E.-M. Krammer, and B. André. Function and regulation of fungal amino acid transporters: Insights from predicted structure. Yeast Membrane Transport, pages 69–106, 2016.

C. H. Graham, D. Storch, and A. Machac. Phylogenetic scale in ecology and evolution. Global Ecology and Biogeography, 27(2):175–187, 2018.

K. Harris and R. Nielsen. The genetic cost of neanderthal introgression. Genetics, 203(2):881–891, 2016.

M. S. Hibbins and M. W. Hahn. Phylogenomic approaches to detecting and characterizing introgression. Genetics, 220(2):iyab173, 2022.

P. M. Hime, A. R. Lemmon, E. C. M. Lemmon, E. Prendini, J. M. Brown, R. C. Thomson, J. D. Kratovil, B. P. Noonan, R. A. Pyron, P. L. Peloso, et al. Phylogenomics reveals ancient gene tree discordance in the amphibian tree of life. Systematic Biology, 70(1):49–66, 2021.

J. L. Humann, T. Lee, S. Ficklin, and D. Main. Gene Prediction: Methods and Protocols. chapter “Structural and functional annotation of eukaryotic genomes with GenSAS”, pages 29–51. Springer, 2019.

M. A. Jarzyna, I. Quintero, and W. Jetz. Global functional and phylogenetic structure of avian assemblages across elevation and latitude. Ecology Letters, 24(2):196–207, 2021.

L. Jiang, J. Tan, and Z. Pu. An experimental test of Darwin’s naturalization hypothesis. The American Naturalist, 175(4):415–423, 2010.

M. Karageorgi, S. C. Groen, F. Sumbul, J. N. Pelaez, K. I. Verster, J. M. Aguilar, A. P. Hastings, S. L. Bernstein, T. Matsunaga, M. Astourian, et al. Genome editing retraces the evolution of toxin resistance in the monarch butterfly. Nature, 574(7778):409–412, 2019.

K. Katoh and D. M. Standley. MAFFT multiple sequence alignment software version 7: improvements in performance and usability. Molecular Biology and Evolution, 30(4):772–780, 2013.

S. N. Kivlin, G. C. Winston, M. L. Goulden, and K. K. Treseder. Environmental filtering affects soil fungal community composition more than dispersal limitation at regional scales. Fungal Ecology, 12:14–25, 2014.

A. M. Kozlov, D. Darriba, T. Flouri, B. Morel, and A. Stamatakis. RAxML-NG: a fast, scalable and user-friendly tool for maximum likelihood phylogenetic inference. Bioinformatics, 35(21):4453–4455, 2019.

S. Lang, F. Erdmann, M. Jung, R. Wagner, A. Cavalie, and R. Zimmermann. Sec61 complexes form ubiquitous ER Ca2+ leak channels. Channels, 5(3):228–235, 2011.

S. Q. Le and O. Gascuel. An improved general amino acid replacement matrix. Molecular biology and evolution, 25 (7):1307–1320, 2008.

J. Liu, J. Huang, Y. Zhao, H. Liu, D. Wang, J. Yang, W. Zhao, I. A. Taylor, and Y.-L. Peng. Structural basis of dna recognition by pcg2 reveals a novel dna binding mode for winged helix-turn-helix domains. Nucleic acids research, 43(2):1231–1240, 2015.

J. B. Losos. Phylogenetic perspectives on community ecology. Ecology, 77(5):1344–1354, 1996.

M. C. Maher and R. D. Hernandez. Rock, paper, scissors: harnessing complementarity in ortholog detection methods improves comparative genomic inference. G3: Genes, Genomes, Genetics, 5(4):629–638, 2015.

U. H. Maier-Greiner, B. Obermaier-Skrobranek, L. M. Estermaier, W. Kammerloher, C. Freund, C. Wülfing, U. I. Burkert, D. H. Matern, M. Breuer, and M. Eulitz. Isolation and properties of a nitrile hydratase from the soil fungus myrothecium verrucaria that is highly specific for the fertilizer cyanamide and cloning of its gene. Proceedings of the National Academy of Sciences, 88(10):4260–4264, 1991.

M. M. Mayfield and J. M. Levine. Opposing effects of competitive exclusion on the phylogenetic structure of communities. Ecology Letters, 13(9):1085–1093, 2010.

O. Meleshko, M. D. Martin, T. S. Korneliussen, C. Schröck, P. Lamkowski, J. Schmutz, A. Healey, B. T. Piatkowski, A. J. Shaw, D. J. Weston, et al. Extensive genome-wide phylogenetic discordance is due to incomplete lineage sorting and not ongoing introgression in a rapidly radiated bryophyte genus. Molecular Biology and Evolution, 38 (7):2750–2766, 2021.

P. Mier, A. J. Pérez-Pulido, E. G. Reynaud, and M. A. Andrade-Navarro. Reading the evolution of compartmentalization in the ribosome assembly toolbox: The yrg protein family. Plos one, 12(1):e0169750, 2017.

M. L. L. Motta, R. R. de Melo, L. M. Zanphorlin, C. A. dos Santos, A. P. de Souza, et al. A novel fungal metal-dependent α-l-arabinofuranosidase of family 54 glycoside hydrolase shows expanded substrate specificity. Scientific Reports, 11(1):1–10, 2021.

T. Münkemüller, S. Lavergne, B. Bzeznik, S. Dray, T. Jombart, K. Schiffers, and W. Thuiller. How to measure and test phylogenetic signal. Methods in Ecology and Evolution, 3(4):743–756, 2012.

A. Muszewska, S. Pi lsyk, U. Perlińska-Lenart, and J. S. Kruszewska. Diversity of cell wall related proteins in human pathogenic fungi. Journal of Fungi, 4(1):6, 2017.

V. K. Nagarajan, C. I. Jones, S. F. Newbury, and P. J. Green. XRN 5’→ 3’ exoribonucleases: structure, mechanisms and functions. Biochimica et Biophysica Acta (BBA)-Gene Regulatory Mechanisms, 1829(6-7):590–603, 2013.

A. Narwani, M. A. Alexandrou, T. H. Oakley, I. T. Carroll, and B. J. Cardinale. Experimental evidence that evolutionary relatedness does not affect the ecological mechanisms of coexistence in freshwater green algae. Ecology Letters, 16(11):1373–1381, 2013.

H. Naughton, M. Alexandrou, T. Oakley, and B. Cardinale. Phylogenetic distance does not predict competition in green algal communities. Ecosphere, 6(7):1–19, 2015.

E. Paradis and K. Schliep. ape 5.0: an environment for modern phylogenetics and evolutionary analyses in R. Bioinformatics, 35(3):526–528, 2019.

H.-S. Park, T.-Y. Nam, K.-H. Han, S. C. Kim, and J.-H. Yu. Velc positively controls sexual development in aspergillus nidulans. PloS One, 9(2):e89883, 2014.

Parmentier, M. Réjou-Méchain, J. Chave, J. Vleminckx, D. W. Thomas, D. Kenfack, G. B. Chuyong, and O. J. Hardy. Prevalence of phylogenetic clustering at multiple scales in an african rain forest tree community. Journal of Ecology, 102(4):1008–1016, 2014.

B. Pease, D. C. Haak, M. W. Hahn, and L. C. Moyle. Phylogenomics reveals three sources of adaptive variation during a rapid radiation. PLoS Biology, 14(2):e1002379, 2016.

G. Peay, M. Belisle, and T. Fukami. Phylogenetic relatedness predicts priority effects in nectar yeast communities. Proceedings of the Royal Society B: Biological Sciences, 279(1729):749–758, 2012.

M. W. Pennell, J. M. Eastman, G. J. Slater, J. W. Brown, J. C. Uyeda, R. G. FitzJohn, M. E. Alfaro, and L. J. Harmon. geiger v2.0: an expanded suite of methods for fitting macroevolutionary models to phylogenetic trees. Bioinformatics, 30(15):2216–2218, 2014.

J. Pöhlmann, C. Risse, C. Seidel, T. Pohlmann, V. Jakopec, E. Walla, P. Ramrath, N. Takeshita, S. Baumann, M. Feldbrügge, et al. The Vip1 inositol polyphosphate kinase family regulates polarized growth and modulates the microtubule cytoskeleton in fungi. PLoS Genetics, 10(9):e1004586, 2014.

M. N. Price, P. S. Dehal, and A. P. Arkin. FastTree 2–approximately maximum-likelihood trees for large alignments. PloS One, 5(3):e9490, 2010.

S. Priebe, C. Kreisel, F. Horn, R. Guthke, and J. Linde. FungiFun2: a comprehensive online resource for systematic analysis of gene lists from fungal species. Bioinformatics, 31(3):445–446, 2015.

R. Ramanujam and N. I. Naqvi. PdeH, a high-affinity cAMP phosphodiesterase, is a key regulator of asexual and pathogenic differentiation in Magnaporthe oryzae. PLoS Pathogens, 6(5):e1000897, 2010.

D. F. Robinson and L. R. Foulds. Comparison of phylogenetic trees. Mathematical Biosciences, 53(1-2):131–147, 1981.

F. J. Rohlf. Comparative methods for the analysis of continuous variables: geometric interpretations. Evolution, 55 (11):2143–2160, 2001.

A. Roy, A. Kumar, D. Baruah, and R. Tamuli. Calcium signaling is involved in diverse cellular processes in fungi. Mycology, 12(1):10–24, 2021.

M. Stanke, R. Steinkamp, S. Waack, and B. Morgenstern. AUGUSTUS: a web server for gene finding in eukaryotes. Nucleic Acids Research, 32(uppl 2):W309–W312, 2004.

S. W. Stevens and J. Abelson. Purification of the yeast U4/U6 U5 small nuclear ribonucleoprotein particle and identification of its proteins. Proceedings of the National Academy of Sciences, 96(13):7226–7231, 1999.

T. Stuwe, C. J. Bley, K. Thierbach, S. Petrovic, S. Schilbach, D. J. Mayo, T. Perriches, E. J. Rundlet, Y. E. Jeon, L. N. Collins, et al. Architecture of the fungal nuclear pore inner ring complex. Science, 350(6256):56–64, 2015.

J. Sukumaran and M. T. Holder. DendroPy: a python library for phylogenetic computing. Bioinformatics, 26(12): 1569–1571, 2010.

N. Taheri-Talesh, T. Horio, L. Araujo-Bazãn, X. Dou, E. A. Espeso, M. A. Penalva, S. A. Osmani, and B. R. Oakley. The tip growth apparatus of aspergillus nidulans. Molecular biology of the cell, 19(4):1439–1449, 2008.

R. Tamuli, R. Kumar, and R. Deka. Cellular roles of neuronal calcium sensor-1 and calcium/calmodulin-dependent kinases in fungi. Journal of Basic Microbiology, 51(2):120–128, 2011.

J. Tannous, C. M. Cosetta, M. T. Drott, T. A. Rush, P. E. Abraham, R. J. Giannone, N. P. Keller, and B. E. Wolfe. Laea-regulated fungal traits mediate bacterial community assembly. mBio, pages e00769–23, 2023.

T. Vasanthakumar and J. L. Rubinstein. Structure and roles of v-type atpases. Trends in biochemical sciences, 45(4): 295–307, 2020.

N. M. Vega and J. Gore. Simple organizing principles in microbial communities. Current opinion in microbiology, 45: 195–202, 2018.

C. O. Webb, D. D. Ackerly, M. A. McPeek, and M. J. Donoghue. Phylogenies and community ecology. Annual Review of Ecology and Systematics, 33(1):475–505, 2002.

B. E. Wolfe, J. E. Button, M. Santarelli, and R. J. Dutton. Cheese rind communities provide tractable systems for in situ and in vitro studies of microbial diversity. Cell, 158(2):422–433, 2014.

M. Wu, J. L. Kostyun, M. W. Hahn, and L. C. Moyle. Dissecting the basis of novel trait evolution in a radiation with widespread phylogenetic discordance. Molecular Ecology, 27(16):3301–3316, 2018.

S. Xu, L. Li, X. Luo, M. Chen, W. Tang, L. Zhan, Z. Dai, T. T. Lam, Y. Guan, and G. Yu. ggtree: a serialized data object for visualization of a phylogenetic tree and annotation data. IMeta, 1(4):e56, 2022.

